# Rapid Ensemble Measurement of Protein Diffusion, Probe Blinking and Photobleaching Dynamics in the Complex Cellular Space

**DOI:** 10.1101/2021.06.22.449491

**Authors:** Simon Sehayek, Xiyu Yi, Shimon Weiss, Paul W. Wiseman

**Author notes:** The work from X. Yi was performed under the auspices of the U.S. Department of Energy by Lawrence Livermore National Laboratory under Contract DE-AC52-07NA27344. Release number: LLNL-JRNL-816054.

## Abstract

We present a fluorescence fluctuation image correlation analysis method that can rapidly and simultaneously measure the diffusion coefficient, photoblinking rates, and fraction of diffusing particles of fluorescent molecules in cells. Unlike other image correlation techniques, we demonstrated that our method could be applied irrespective of a non-uniformly distributed, immobile blinking fluorophore population. This allows us to measure blinking and transport dynamics in complex cell morphologies, a benefit for a range of super-resolution fluorescence imaging approaches that rely on probe emission blinking. Furthermore, we showed that our technique could be applied without directly accounting for photobleaching. We successfully employed our technique on several simulations with realistic EMCCD noise and photobleaching models, as well as on Dronpa-C12 labeled beta-actin in living NIH/3T3 and HeLa cells. We found that the diffusion coefficients measured using our method were consistent with previous literature values. We further found that photoblinking rates measured in the live HeLa cells varied as expected with changing excitation power.

## INTRODUCTION

The past decade has seen a revolution in optical microscopy with the advent of far-field super-resolution approaches. Fluorescence super-resolution microscopy has become an invaluable tool for furthering our understanding of biological systems by allowing one to circumvent the diffraction limited resolution of traditional fluorescence microscopy techniques, leading to important insights in cell biology, neuroscience and cellular biophysics.[1–5] Among the super-resolution imaging techniques, single-molecule localization microscopy (SMLM) methods are one of the most commonly used. These methods rely on photophysical processes to localize the position of single fluorophores with a spatial uncertainty much lower than the wavelength diffraction limit of light. Some popular examples of such techniques are stochastic optical reconstruction microscopy (STORM)[6] and photoactivated localization microscopy (PALM).[7] Super-resolution optical fluctuation imaging (SOFI)[8] also relies on the stochastic photoswitching nature of fluorophores, but builds super-resolution images using the cumulants of the fluorescence fluctuations. Many applications of super-resolution, however, have so far been limited to studying immobile components in cells, or static molecules in chemically fixed cells, without examining their dynamic counterparts. While super-resolution has also been coupled with quantitative methods to investigate these dynamics, there are currently only a limited number of approaches that combine super-resolution imaging and measurement of dynamics. Notably, stimulated emission depletion (STED) microscopy was combined with fluorescence correlation spectroscopy (FCS) in STED-FCS,[9] which was shown to better characterize heterogeneous diffusive behavior of membrane biomolecules than traditional FCS by reducing the beam spot size. Scanning versions of this technique exist, as well.[10, 11] Single-particle tracking (SPT) and PALM were also combined in sptPALM.[12] Minimal photon fluxes (MINFLUX)[13] is also capable of super-resolved SPT. Another example is fcsSOFI, which combines FCS and SOFI techniques to form super-resolved diffusion maps.[14]

Many methods currently exist for measuring the dynamics in biological systems and a significant subset rely on measurement of fluorescence fluctuations to compute autocorrelations that are then fit with specific models to measure transport parameters, such as biomolecule diffusion coefficients and directed flow rates. Conventional FCS[15–17] is a widely used example that analyzes fluorescence fluctuation time series collected from a single fixed laser focal spot. Imaging FCS[18] was developed to allow for multiplexed FCS analysis of pixels forming a fluorescence image. Similarly, image correlation spectroscopy (ICS) methods[19–22] also analyze fluorescence fluctuations in images, but utilize both spatial and temporal information when computing the autocorrelation. One advantage of ICS techniques is that the use of spatial information increases statistical sampling; however, the inherent spatiotemporal heterogeneity in biological systems makes it difficult to simply average over different pixels and frames. ICS techniques usually handle spatiotemporal heterogeneity by analyzing many smaller local regions of interest (ROIs), and then correlating over a chosen time of interest (TOI). This is the approach for generating flow maps when using spatiotemporal image correlation spectroscopy (STICS).[21] The most challenging example of heterogeneity in time in fluorescence microscopy is the issue of photobleaching, which has been addressed using both pre-[23–25] and post-processing[26–30] methods. Another example is anomalous diffusion, which has been investigated in multiple FCS studies.[31–36]

As super-resolution aims to better resolve complex cell processes in space and time, a technique that can analyze dynamics in the presence of spatially and temporally heterogeneous structures is required. Here we present an image correlation method that can successfully and rapidly analyze heterogeneous systems for accurate measurement of biomolecule diffusion coefficients, probe photoblinking rates and fraction of particles undergoing diffusion (see Figure 1). Along with measuring dynamic parameters in a complex cell environment, we anticipate that the measurement of probe photoblinking rates will be useful for optimal fluorescent probe development and optimization for methods like STORM and SOFI. The ability of our method to analyze spatially heterogeneous systems further allows us to significantly increase the spatial sampling used in our autocorrelation computation.

**FIG. 1:**
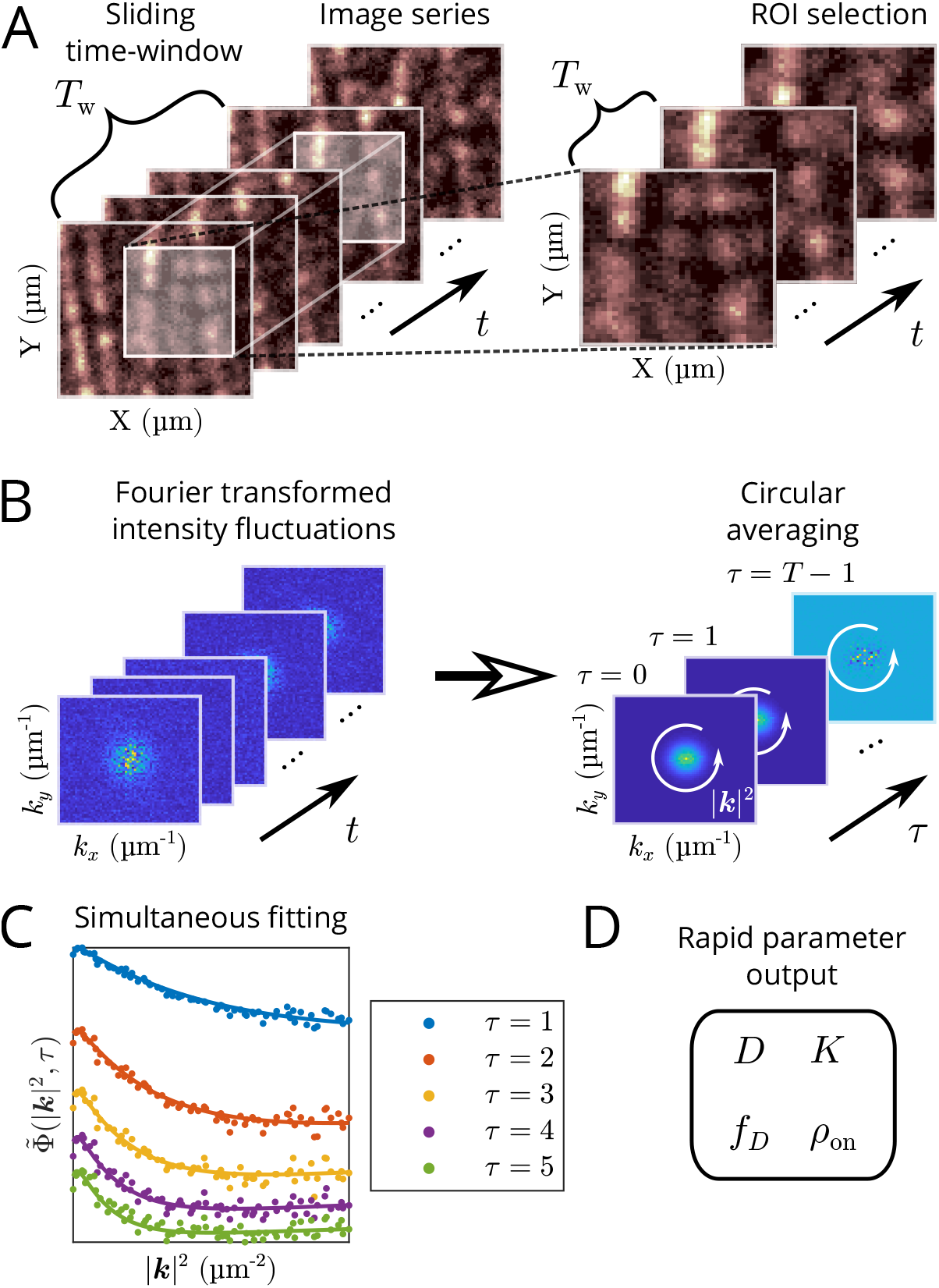
Schematic illustrating extended kICS method application. (A) An ROI and sliding time-window are first chosen from an image series. Intensity fluctuations are computed locally in time, according to the chosen time-window size, to mitigate the effects of photobleaching. (B) The 2D spatial Fourier transforms of the intensity fluctuations are first computed in each frame, then this k-space ROI frame stack is autocorrelated in time. A circular averaging of the autocorrelation is also calculated when the dynamics of the system are isotropic. Non-uniformly distributed, immobile blinking fluorophore populations do not systematically affect the fluctuation defined autocorrelation. (C) Computed ACF (points) and simultaneous fits (lines) over five time-lags. (D) The process of computing the ACF and fitting is rapid (order of seconds for 64 × 64 pixel image series with 2048 frames) and outputs: the diffusion coefficient (*D*), the sum of the photoblinking rates (*K*), the fraction of time spent in the on-state (*ρ*_on_), and the fraction of diffusing particles relative to all particles (*f_D_*).

Our method is based on k-space image correlation spectroscopy (kICS),[19] which was originally developed for measuring transport parameters independently from the fluorophore photophysics. Similar to other ICS techniques, one of the underlying assumptions behind kICS is uniformity in space and time in the data being analyzed.[20] Non-uniformity in space caused by the presence of immobile structures were commonly filtered out by either subtracting the time average, or equivalently, applying a Fourier filter. This solution does not work, however, when the fluorophores labeling these structures are undergoing photophysical processes *e*.*g*. photoblinking and photobleaching on timescales comparable to the imaging, thus making them difficult to analyze with traditional ICS methods. This is especially relevant in the context of SMLM and SOFI methods. Therefore, this significant extension of kICS is important for dealing with real heterogenities in space and time in the complex cellular environment.

We begin by showing that our definition of the autocorrelation is approximately independent of immobile particle positions. We then derive the autocorrelation for a mixed system of photoblinking immobile and diffusing particles that is independent of any direct imaging parameters in 2D, such as the point spread function (PSF). We proceed to demonstrate the accurate measurement of diffusion and photophysical parameters, when applying our technique on simulations. We then employ our method on Dronpa-labeled beta-actin (*β*-actin) imaged in live 3T3 and HeLa cells. Our measured diffusion coefficients for *β*-actin from these experiments are consistent with previously reported values. Furthermore, we observed that the photoblinking follows the expected dependence on excitation power in our HeLa cell experiments.

## THEORY

In the development of the original kICS technique by Kolin et al. (2006),[19] it was shown that diffusion and flow dynamics could be recovered regardless of the photophysical properties of the flurophores (*e*.*g*. photoblinking and photobleaching). However, the original method did not consider the presence of immobile particle populations also undergoing photophysical processes. We show here that with the presence of such populations, one cannot simply apply the original kICS analysis technique. We further derive an expression for the autocorrelation function (ACF) that we simultaneously fit for transport and photo-physical parameters. This expression is independent of any direct imaging parameters in 2D (*e*.*g*. PSF size, amplitude, *etc*.), as well as the immobile particle positions, allowing us to probe systems with complex immobile particle spatial arrangements and non-uniform cell morphologies. Furthermore, this enables us to maximize statistical spatial sampling when computing the ACF. This is in contrast with other image correlation techniques, which require spatially uniform regions for analysis.[20] Consequently, previous image correlation techniques would not be able to include cell boundaries and narrow projections, such as dendritic spines and narrow cellular lamellae, such as dendritic projections, when choosing ROIs for analysis. The 2D model presented here can be used to model membrane dynamics sampled using TIRF microscopy, for example. We later extend this to the 3D case when analyzing the live 3T3 and HeLa cell data. We will refer to the developed technique as an extended kICS method.

We begin with the definition of a fluorescence microscopy image series of intensities in 2D, *i*(***r***, *t*):

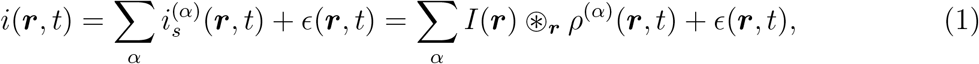

where 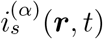 is the fluorescence intensity from the labeled particles at position ***r*** and time *t*, belonging to population *α* with common transport parameters; *I*(***r***) is the PSF; ⊛_***r***_ is a spatial convolution; *ϵ*(***r***, *t*) is an additive noise term, which is assumed to be independent from itself for any (***r***, *t*) ≠ (***r***′, *t*′); and *ρ*^(*α*)^(***r***, *t*) is the apparent particle density of population *α*, *i.e.*, the density of particles in population *α* that are emitting and detectable. Note we only consider dependence of population *α* on particle position in this work. The apparent density of particles belonging to population *α* is given by:

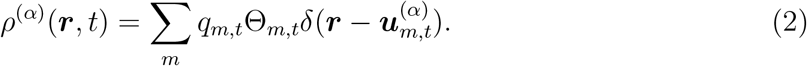

In this last equation, *δ*(·) is the 2-dimensional Dirac delta function; *m* is an index denoting the fluorophores (which we also refer to as “particles”); *q_m,t_* is the instantaneous rate of detector counts for the *m*^th^ fluorophore at time *t*, which depends on several factors, including the photon budget, quantum efficiency of the detector, and camera detector gain; 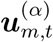 is the position of the *m*^th^ fluorophore at time *t* (belonging to population *α*), and Θ*_m,t_* is its photo-emissive state, expressed as:

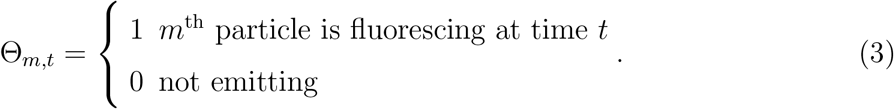

Note we do not consider photophysical transitions between singlet states (*i.e.*, absorption/emission and quenching), as these are far beyond the time-resolution capabilities of electron multiplying charge-coupled device (EMCCD) camera detectors.

Since we developed this approach for widefield fluorescence microscopy, we account for faster processes by also considering the effects of camera detector integration time by substituting:[30]

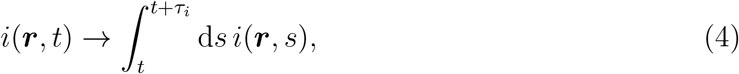

with *τ_i_* being the integration time (in this work, we consider *τ_i_* = 1); however, we leave the integration out of the notation, as it is straightforward to redo the derivation with it.

Defining the spatial Fourier transform as 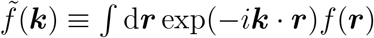, we have the Fourier counterparts of Eqs. (1) and (2); respectively,

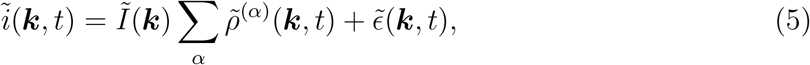

and,

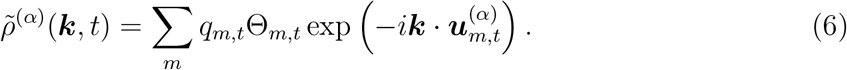

In this work, we calculate the autocorrelation as:

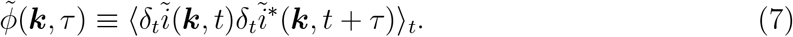

Practically, the above autocorrelation is realized by computing a time average, therefore we use the notation ⟨…⟩*_t_* to denote an expectation value that only considers random variables that depend on time to be random. We also define *δ_t_* as a fluctuation with respect to the time average, *i.e.*,

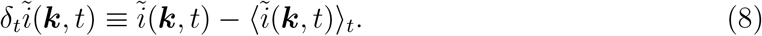

Note in the original kICS work,[19] the autocorrelation was defined without the fluctuations. In the Supporting Information (SI), we show that the original definition leads to noisy autocorrelations affected by the immobile blinking particle positions (see SI:Comparison with original kICS method). Using the Fourier transform of an image series in Eq. (5), we can express the autocorrelation in Eq. (7) as:

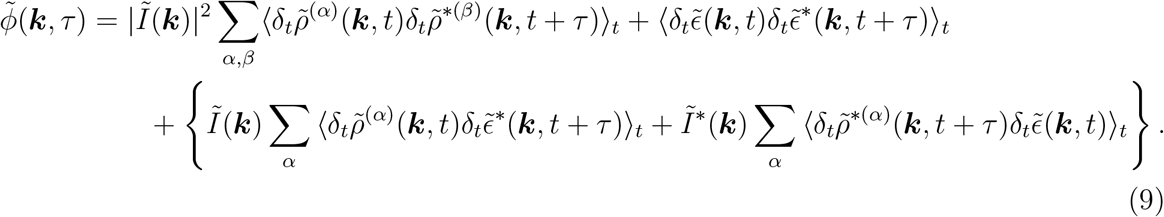

The terms in the curly brackets are zero, assuming noise and particle positions are independent. We show in SI:Noise autocorrelation that the autocorrelation of the noise (*i.e.*, the second term) only affects *τ* = 0 by a constant offset, which we will denote by 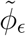. Note |***k***|^2^ = 0 is also affected by the noise, but is omitted from the analysis.

Using Eq. (6), for immobile populations we have:

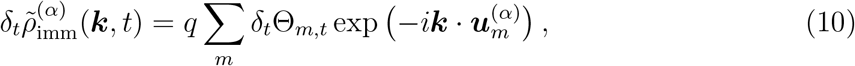

where we have assumed all fluorophores have equal quantal brightness (*i.e.*, *q_m,t_* = *q*). Note further that the time indexing is dropped for immobile particle positions so that they are unaffected by the time averaging. Conversely, for mobile populations:

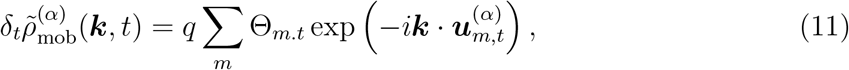

since 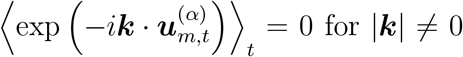, assuming the mobile particle positions are uniformly distributed in space within the chosen ROI.

Using Eqs. (10) and (11), Eq. (9) for one mobile and one immobile population becomes:

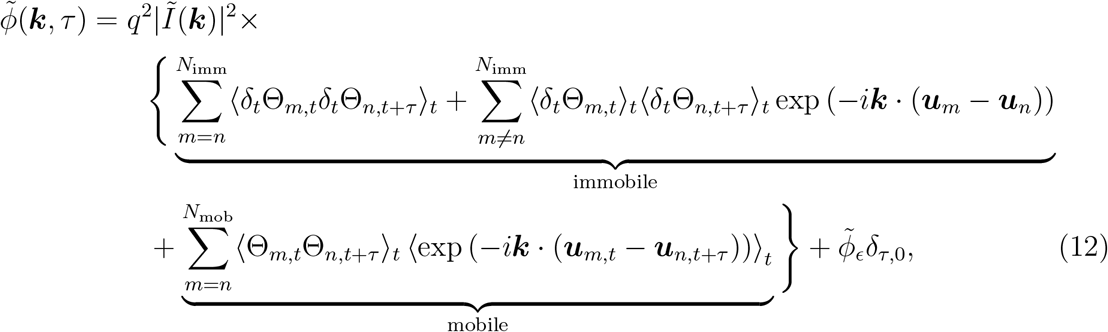

where we have assumed non-identical particles are mutually independent, so that we can sum their individual autocorrelations, and *δ*_*τ*,0_ is the Kronecker delta function. Note the fluctuations are not necessarily expected to vanish in the second term since, in practice, they are computed by subtracting the sample time average, which does not converge to the ensemble average in non-ergodic systems. Furthermore, subtraction by the spatial average only affects the autocorrelation at |***k***| = 0. Notice also the second term is affected by the immobile particle positions, which is why we need to define conditions when it is approximately zero. It is clear that without photobleaching (or any other non-stationary photophysical process) the photophysical fluctuations are indeed zero on average; however, in the presence of bleaching the fluctuations need to be properly defined to make sure the second term in Eq. (12) is approximately zero. For this reason, we use *local* time-averaging over a subset time-window to compute Eq. (8) in practice (see “Autocorrelation computation” in Materials and Methods), *i.e.*,

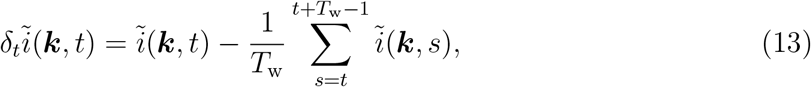

where *T*_w_ is the window size chosen for the local time average. If the photobleaching is slow and *T*_w_ is large enough, then Eq. (8) is a good approximation of Eq. (13). On the other hand, if the photobleaching is significant one must account for *T*_w_ in the autocorrelation (see SI: Time-windowed correction, where we provide an expression for this correction). Eq. (8) will also not hold when the dynamics in the system are slow and resemble immobility; in this case, the effects of time-windowing should again be accounted for (for example, see Figure 2 (C)). Note that if the second term in Eq. (12) is zero, then the autocorrelation is independent of any assumptions on immobile particle positions, making it a powerful tool to study previously inaccessible systems using image correlation. Time-windowed or moving-average subtraction has been previously used in raster ICS (RICS) as a way to filter out slow-moving fluorescent objects.[22, 37]

In order to obtain a quantity that is independent of PSF, we define the autocorrelation function (ACF) as:

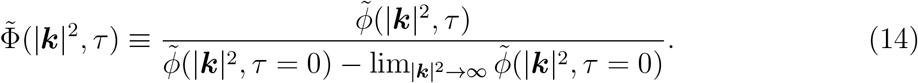

**FIG. 2:**
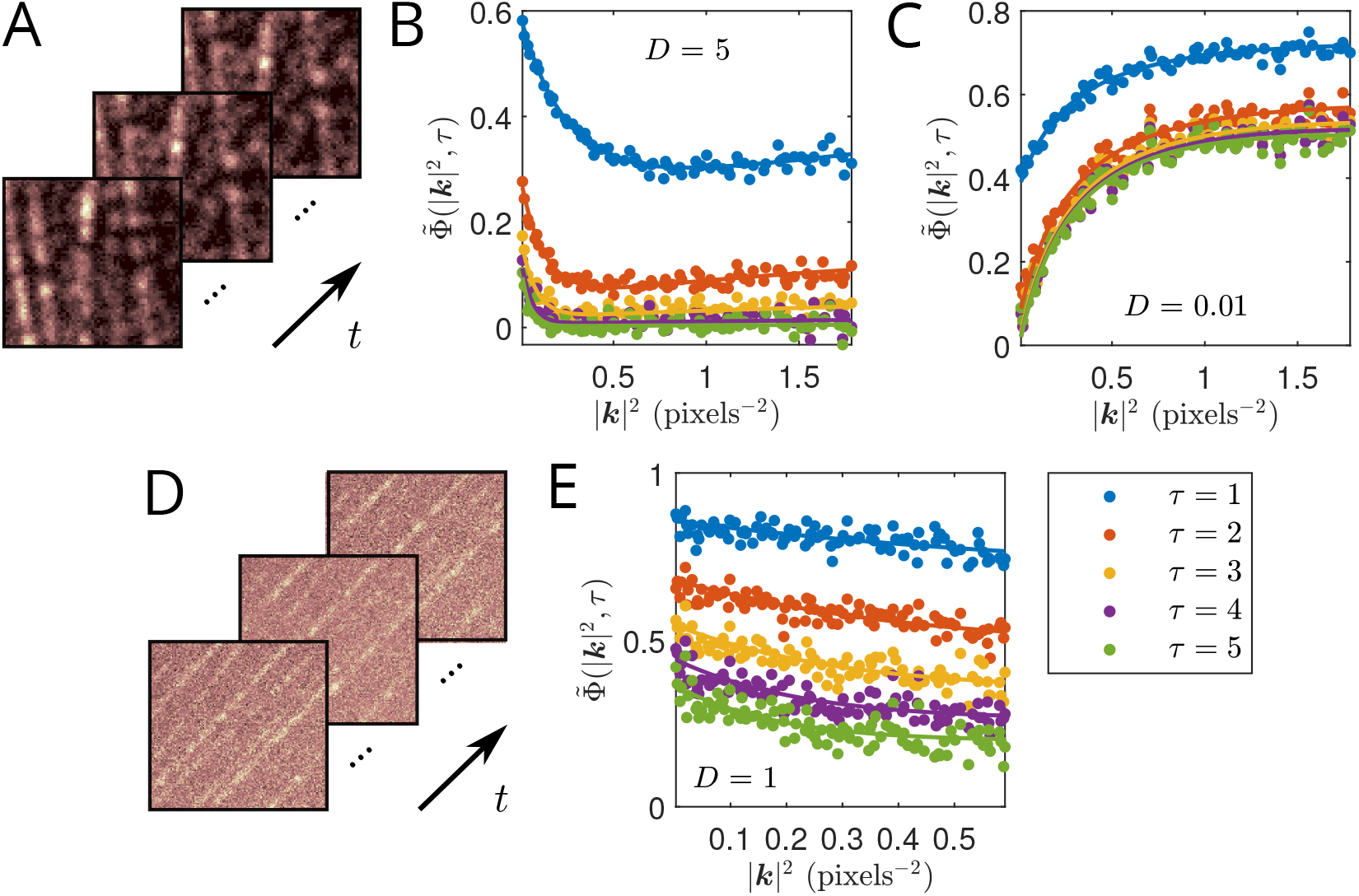
Example ACFs and fits computed from simulations of filamentous structures composed of static blinking particles, with a second population of freely diffusing particles. Both populations are set to have the same photophysical properties. Each simulation also contained simulated EMCCD noise. The ACF fits were all done over the first 10 time-lags; only the first 5 are shown. Time windowing was done with *T*_w_ = 200 frames in each case. Simulation and fit details are shown in Table I. (A) Example of simulated intensity images in time. (B, C) Computed ACFs (points) and corresponding simultaneous fits (lines) for simulations with *D* = 5 and 0.01 pixels^2^/frame, respectively. In (C), we utilize a fit model that corrects for the chosen sliding time-window. (D) Sample simulation with higher simulated autofluorescence background and lower photon budget than simulation shown in (A). (E) Corresponding computed ACF and fit of simulation shown in (D) with *D* = 1 pixels^2^/frame.

**TABLE I:**
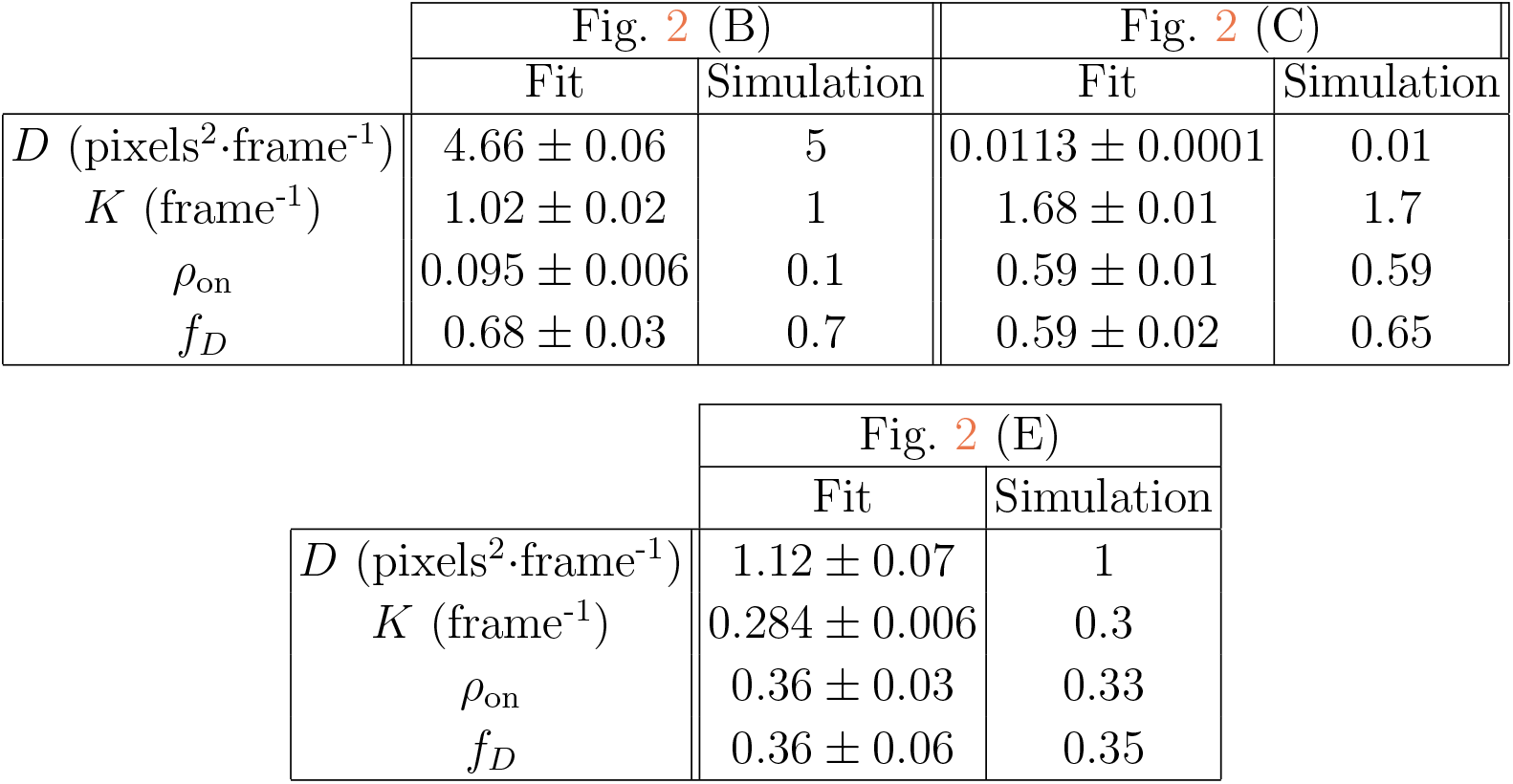
Comparison of fitted and simulated parameters for fits shown in Figure 2.

The second term in the denominator removes dependence of the ACF on the noise; practically, it is computed as the large |***k***|^2^ offset in the autocorrelation. Furthermore, if the mobile components of the system being analyzed are isotropic, one can circularly average the autocorrelations for statistical sampling purposes.[19, 20, 38] Since the focus of this work will be on such systems, the dependence of the ACF in this last equation is left to be on |***k***|^2^. It should also be mentioned that this definition of the ACF leads to division by zero after sufficient decay of the PSF, so that the ACF must be appropriately trimmed in |***k***|^2^ prior to fitting. For a 2D mixture of diffusing and immobile particles, Eq. (14) can be written explicitly as:[19, 30, 39]

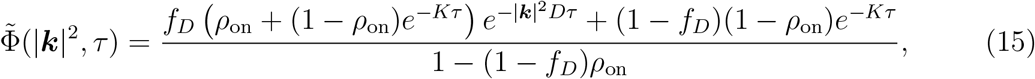

where *D* is the diffusion coefficient; *K* ≡ *k*_on_+*k*_off_ is the sum of the photoblinking rates, where *k*_on_ and *k*_off_ are the on- and off-blinking rates, respectively; *ρ*_on_ is the fraction of time spent in the fluorescent on-state; and *f_D_* is the fraction of particles diffusing. Note that the blinking in this last expression is assumed to follow a two-state on-off model, without photobleaching. In the SI, we include a derivation considering the effects of bleaching and detector time-integration, as well as a derivation for the 3D effects on the ACF (see SI:Autocorrelation function derivation).

## RESULTS AND DISCUSSION

### Computer simulations

We first apply our extended kICS method on 2D computer simulations of immobile filamentous structures composed of fixed emitting particles, with a second simulated population of freely diffusing particles, as shown in Figure 2. The fit results from this figure are tabulated in Table I. The fitted parameters demonstrate the wide range of diffusion coefficients that are measurable in heterogeneous morphologies using our extended technique.

The simulations assumed both immobile and diffusing particle populations to have the same photoblinking and photobleaching rates. Each simulation also contained simulated EMCCD noise (see Sehayek *et al.* (2019)[30] for noise model details). Note the non-uniformity in the immobile particle positions placed along the simulated filaments in Figure 2, which confirms that the technique can be successfully applied in non-homogeneous systems. Conversely, previous image correlation techniques required ROIs to be selected where the spatial distribution of particles was homogeneous *e*.*g*. avoiding cell boundaries (see SI:Comparison with original kICS method where we compare our extended kICS technique to the original one).[20] It would be impossible to select such an ROI with both diffusing and immobile particles in the simulations shown in Figure 2. The ability to analyze larger ROIs further enables us to increase the spatial sampling in our analyses; however, we assume that all the particles within this ROI have transport and photophysical parameters that are drawn from common distributions, *i.e.*, we assume a single diffusing population and a single photophysical population (while the extension to multiple populations with different dynamic parameters is possible, one must be mindful of the possibility of overfitting). Thus, our analysis offers a coarse-grained approach for quickly measuring these parameters within regions. SPT is beneficial when a common distribution cannot be assumed for such parameters. However, the fluorescence correlation approach can be applied to cell expression systems where high density labeling might not permit SPT. SPT is also limited by factors such as photoblinking in transport populations.

In Figure 2 (B), the shape of the ACF is characterized by a decay at low |***k***|^2^, and a convergence to a non-zero value at higher |***k***|^2^. The former is attributed to the diffusion coefficient, while the latter is due to the presence of an immobile particle population, as can be seen from Eq. (15) (note that these fits use the time-integrated version of this equation; see SI:Autocorrelation function derivation for details). The decrease in the ACF amplitude and offset with increasing time-lag is due to the photophysical processes in the system, *i.e.*, photoblinking and photobleaching.

When the ACF is characterized by an initial increase along |***k***|^2^, as is the case in Figure 2 (C), one needs to account for the effect of the sliding time-window (see SI:Time-windowed correction). This type of behavior occurs when the diffusion is relatively slow, or when the chosen time-window is relatively short.

In Figure 2 (D), we show sample images from a simulation with more noise. Specifically, we increased the simulated autofluorescence background, while decreasing the photon budget of the simulated fluorophores. To accurately analyze such an image series (Figure 2 (E)), we required larger ROIs than the ones used in the previous analyses. Consequently, we generated this simulation on a 256 × 256 pixel grid.

Note that photobleaching was not considered in the fits shown in Table I. A derivation considering the effects of bleaching on the ACF is included in SI:Time-windowed correction. Although one can consider these effects, we have demonstrated that we can still obtain accurate parameters in the presence of photobleaching, without having to incorporate it into our fit model (given appropriate choice of time-window). This is beneficial as bleaching pathways of a fluorescent label are not often known.

We also note that the fraction of time spent in the on-state, *ρ*_on_, cannot be measured for a purely immobile population, *i.e.*, *f_D_* = 0. This can be seen by the lack of dependence on *ρ*_on_ in Eq. (15) in this limit. Previous techniques have described how to measure the photoblinking rates for immobile emitters.[30, 40–43]

To perform our analysis, it is essential to choose a time-window, an ROI, a range of time-lags to fit simultaneously, and cutoff values for |***k***|^2^. As discussed in the Theory section above, the choice of time-window will mainly depend on the photobleaching rate. A value of *T*_w_ that is too small results in loss of information, but a smoother ACF. A value of *T*_w_ that is too large will result in a noisier ACF that is more influenced by the photobleaching, which may lead to poorer fits. With a simulated bleaching rate of *k*_p_ = 10^−4^ frames^−1^ (with a frame acquisition time of 50 ms, this corresponds to a characteristic bleaching time of 500 s), we find a suitable choice of time-window to be *T*_w_ = 200 frames. We will use this value consistently throughout this work for the same value of *k*_p_. The same value of *T*_w_ can be used for smaller bleaching rates. In general, we found from our simulations that *T*_w_ can be chosen to be about 2–5% of the characteristic bleaching time.

As mentioned above, when choosing an ROI it is necessary to select a relatively large region, in order to avoid aliasing of the ACF along |***k***|^2^. This is especially important when dealing with larger diffusion coefficients, as the decay will appear in a short range of small |***k***|^2^.

Similarly, when choosing a range of time-lags to fit, it is best to choose a wide range, in order to capture slower dynamics. If the range is too large, the simultaneous fitting will be visibly biased and a smaller range should then be used. Trying to fit a larger time-lag range can be complicated by the presence of photobleaching, for example (we show how to account for these effects in SI:Time-windowed correction). We also remark that the ACF is noisier for higher time-lags, so that it is informative to compare them with their fits to gauge whether the fitted time-lag range is too wide. We showed that we can achieve reasonable fits for our simulated parameters by fitting the 10 first time-lags in our analyses in Table I. Systematic errors were reduced when including more time-lags in the fit, or when choosing larger ROIs, but in non-simulated data, it may be difficult to make such adjustments.

Finally, we discuss choosing cutoff values for |***k***|^2^. We discard the value of the ACF at |***k***|^2^ = 0 since it is affected by the noise in the system, as was mentioned in the Theory section. Excluding a few small |***k***|^2^ is also beneficial for avoiding the autocorrelation from the time-windowing, when not using the time-window correction (see SI:Time-windowed correction for details). The maximum value for |***k***|^2^ can be selected by examining where the ACF begins to diverge due to the normalization in Eq. (14). It is optimal to choose the largest possible range of |***k***|^2^ to fit while avoiding points that are too noisy due to the normalization. Choice of the maximum cutoff will depend on PSF size as well as the noise in the system. As was previously mentioned, this is to avoid division by zero that occurs due to our definition of the ACF in Eq. (14). This occurs because the noise is subtracted from the denominator, which is then effectively zero after the PSF has sufficiently decayed. In Figure 2 (E), due to the higher noise in the simulation, the fit has a smaller chosen maximum cutoff of |***k***|^2^ since higher values result in a divergence of the ACF.

Each simulation had photobleaching rate *k*_p_ = 10^−4^ frame^−1^ over *T* = 2048 frames. Simulations (B) and (C) were generated on 128 × 128 pixel grids, while (E) was on a 256 × 256 pixel grid. The simulations were assigned 8130, 8411, 9048 total particles, respectively. In each case, fitted parameters and errors were obtained by splitting the simulation spatially into 4 equally sized and independent ROIs and then calculating the mean and its standard error from their analyses.

In Figure 3, we perform the same analysis on a simulated dendritic morphology. A comparison of the simulated and fitted parameters recovered using our analysis is shown in Table II. This further demonstrates the ability of the technique to be applied independently of immobile particle distribution. In this case, it would again be impossible to select an ROI containing a uniform distribution of both diffusing and immobile populations for analysis using previous image correlation techniques. As mentioned previously, with the extended kICS technique developed in this work, we are no longer restricted by this requirement and can further benefit from large ROIs to increase the spatial sampling in our analysis.

**FIG. 3:**
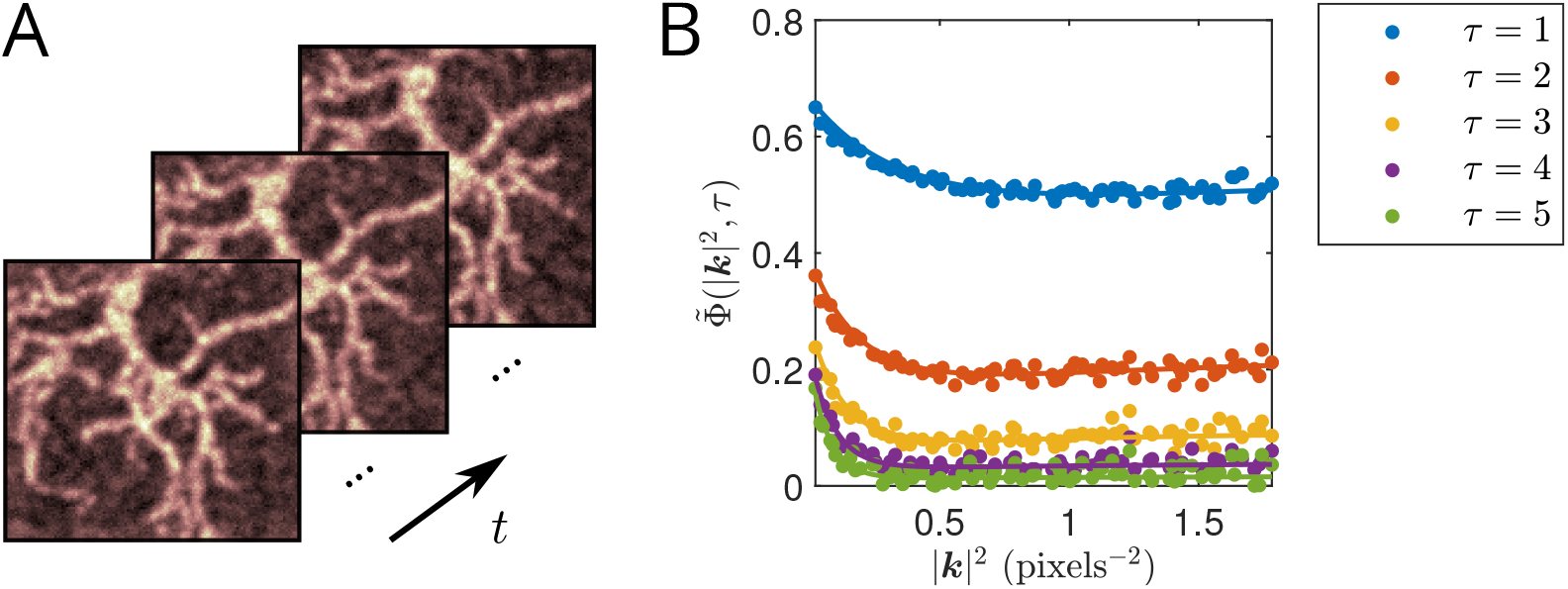
Example ACF computed from simulation of dendritic structures and fit. (A) Sample simulated intensity images in time. Simulations contain immobile particles on the dendrites, along with a background of freely diffusing particles. Both populations are assumed to have equal photophysical properties. (B) Computed ACF (points) and corresponding simultaneous fit (lines). The fit is done over the first 10 time-lags; only first 5 are shown. Time windowing was done with *T*_w_ = 200 frames. Fit details are shown in Table II.

**TABLE II:** Comparison of fitted and simulated parameters for fits shown in Figure 3.

|  | Fit | Simulation |
| --- | --- | --- |
| $D$ (pixels<sup>2</sup>·frame<sup>-1</sup>) | $2.80 \pm 0.08$ | 3 |
| $K$ (frame<sup>-1</sup>) | $0.86 \pm 0.01$ | 0.9 |
| $\rho_{\text{on}}$ | $0.38 \pm 0.03$ | 0.44 |
| $f_D$ | $0.26 \pm 0.02$ | 0.3 |

Simulation had photobleaching rate *k*_p_ = 10^−4^ frame^−1^ and was generated on 128 × 128 pixel grid with 10744 total particles and *T* = 2048 frames. Fitted parameters and errors were obtained by splitting the simulation spatially into 5 64 × 64 ROIs, with some overlap between different parts, and then calculating the mean and its standard error from their analyses. The ROIs were chosen so that a significant portion of the simulated dendritic structure was encompassed, overall.

### Live NIH/3T3 cell data

Using a widefield fluorescence microscope equipped with an EMCCD camera detector, we imaged *β*-actin labled with Dronpa-C12 in an NIH/3T3 fibroblast cell line expression system. The Dronpa-C12 exhibited blinking and photobleaching during image acquisition and the *β*-actin pool was both diffusively mobile within the cell and immobile in actin filaments. A sample of our analysis from the data is shown in Figure 4.

**FIG. 4:**
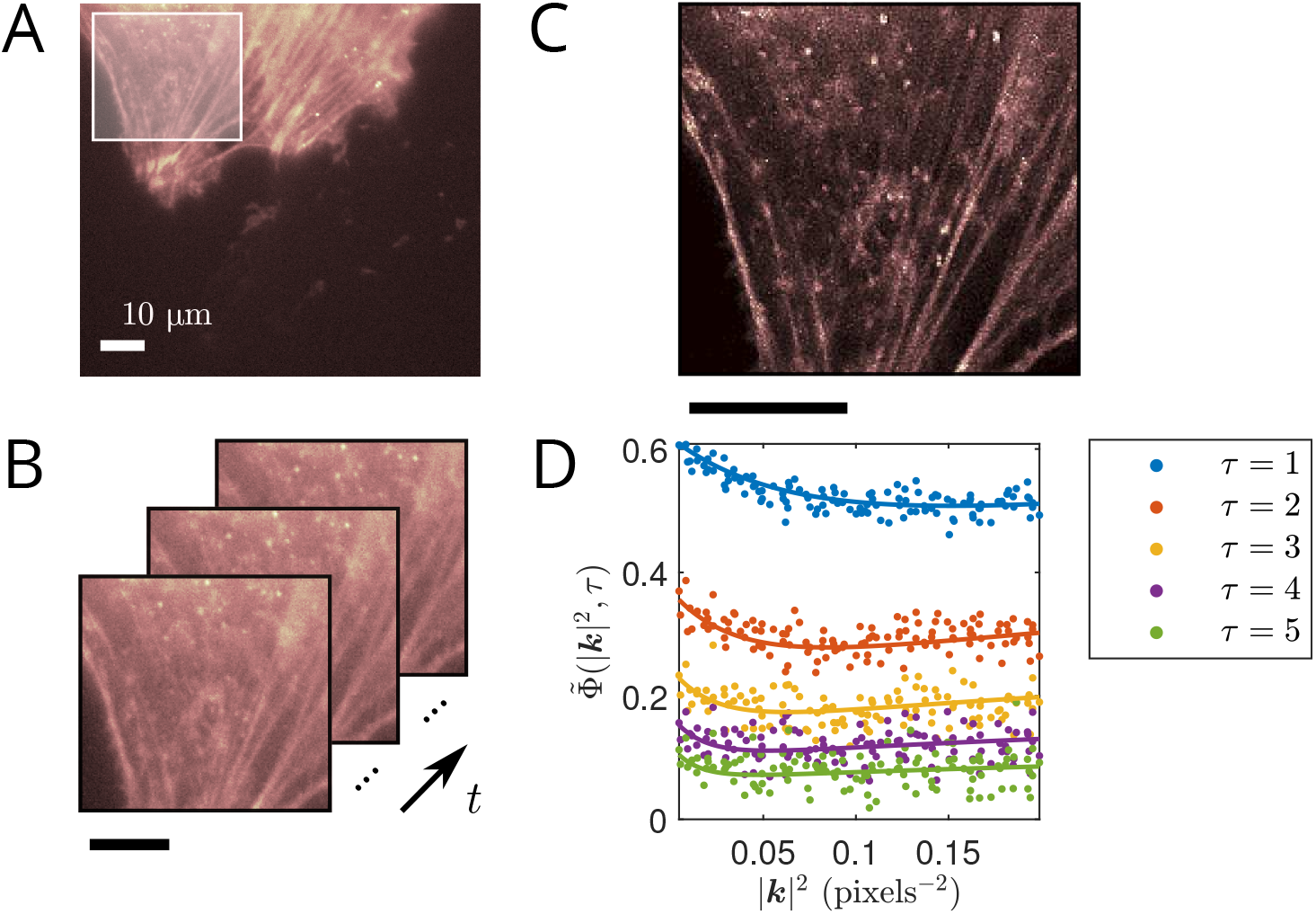
ACF computed from Dronpa-C12 labeled actin in a live NIH/3T3 cell and fit. (A) Fluorescence image of Dronpa-C12 labeled actin in a live NIH/3T3 cell, with ROI used in analysis highlighted. (B) Sample fluorescence images in time of ROI shown in (A). (C) SOFI image generated from immobile blinking fluorophores (see GitHub code for SOFI 2.0). (D) Computed ACF from ROI shown in (A) (points) and corresponding simultaneous fit (lines). The fit was done over the first 5 time-lags. Time-windowing was done with *T*_w_ = 100 frames. Fitted parameters: *D* = 9.2 ± 0.4 μm^2^ · s^−1^, *K* = 7.6 ± 0.4 s^−1^ and *f_D_* = 0.58 ± 0.03. Fit for *ρ*_on_ omitted due to inconsistent values between different ROIs and TOIs. Pixel size: 177.78 μm. Frame time: 50 ms. Analysis details: Two spatially independent ROIs (each about 30 × 30 μm^2^) over two temporally independent TOIs (each about 50 s in length) were considered in the analysis. The reported fitted parameters and errors are given as the mean and its standard error from these analyses.

In Figure 4 (C), we reconfirm that the blinking of immobile fluorophores can be used to obtain a SOFI[8] image using SOFI 2.0[44, 45] (see GitHub code for SOFI 2.0). Along with our dynamic analysis of this data, we demonstrate that we can extract both static and dynamic information from our system with careful selection of correlation analysis approaches.

Since the data was acquired using a widefield fluorescence microscope with actin monomers diffusing in the cytoplasm, we needed to employ a 3D model for the ACF fit (see SI:Diffusing and immobile populations (3D) for more details). The extension to a 3D model (without considering integration time effects) is achieved through a scaling factor that depends on *τ*, and consequently, does not affect the behavior of the ACF along |***k***|^2^. Using the 2D fit model for the ACF yielded visibly inconsistent fits to the data. Notably, our reported value for the apparent diffusion coefficient (9.2 ± 0.4 μm^2^ · s^−1^) is within range of the simulated and experimentally verified diffusion coefficient of globular actin (G-actin) in the cytoplasm of ~ 3–30 μm^2^·s^−1^.[46–50] This is also in reasonable agreement with the diffusion coefficient of Dronpa-labeled actin in an MCF-7 cell of 13.7 μm^2^· s^−1^, reported by Kiuchi *et al.* (2011)[49] Our measurement of a relatively rapid diffusion coefficient is further confirmed by the behavior of the ACF, which exhibits a characteristic initial decay in |***k***|^2^, as in Figure 2 (B). This is in contrast to systems with lower diffusion coefficients, which exhibit an initial increase in |***k***|^2^, as in Figure 2 (C). As can also be seen from Figure 4, our assumption of a single diffusing population provides a reasonable fit to the data. Therefore, we argue that it would not be advantageous to include more diffusing components in our fit model since it would risk overfitting our data.

In order to account for 3D effects, we needed to first estimate the *e*^−2^ radius along the axial direction from the data. To this end, we used the Abbe resolution criterion to determine the full width at half maximum of the PSF in *z*, *i.e.*, 2*λ/*NA^2^, which was then converted to an *e*^−2^ radius.

McGrath *et al.* (1998)[46] demonstrated that they can simultaneously measure actin filament turnover rate, fraction of actin in filaments and actin diffusion using either fluorescence recovery after photobleaching (FRAP) or photoactivation of fluorescence (PAF). In their model, the filamentous actin is not necessarily immobile, but not diffusing. We show that on the time and spatial scales we considered in our analysis, the actin flow is negligible. Furthermore, filament turnover rate is an important parameter at filament ends, but we chose ROIs away from these ends, so that we would not have to consider such effects.

Since flow appears as an imaginary component in the autocorrelation,[19] we compared the magnitude of the imaginary part to that of the modulus of the autocorrelation, *i.e.*,

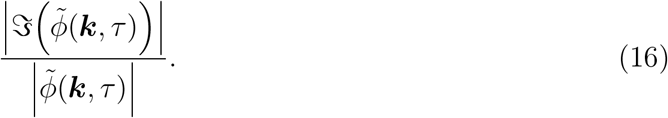

This quantity was determined to be very small (close to machine precision), confirming that the flow is negligible relative to other dynamics over the time and spatial scales examined.

From our analysis of the data shown in Figure 4, we also found a diffusing fraction of *f_D_* = 0.58 ± 0.03. Gasilina *et al.* (2019)[51] reported a percentage of filamentous actin (F-actin) of 48 ± 4% from their immunoblotting-based analysis of wild type NIH/3T3 fibroblast cells. If we assume F-actin to be immobile and G-actin to be diffusing, then this value corresponds to *f_D_* = 0.52 ± 0.04.

We further tested whether the G-actin was undergoing anomalous subdiffusion in the cell. This can approximately be done by replacing the dependence of the ACF from *τ* → *τ^α^* (ignoring detector time integration), where *α* is the degree of subdiffusion.[34] A fit including *α* as a free parameter yielded a fitted value of *α* ~ 1, indicating that the diffusion of the G-actin within this non-migrating cell was mainly free diffusion.

### Live HeLa cell data

We proceeded to analyze Dronpa-C12 labeled *β*-actin in live HeLa cells imaged under different excitation intensities. Our analyses are shown in Figure 5, with results given in Table III below.

**FIG. 5:**
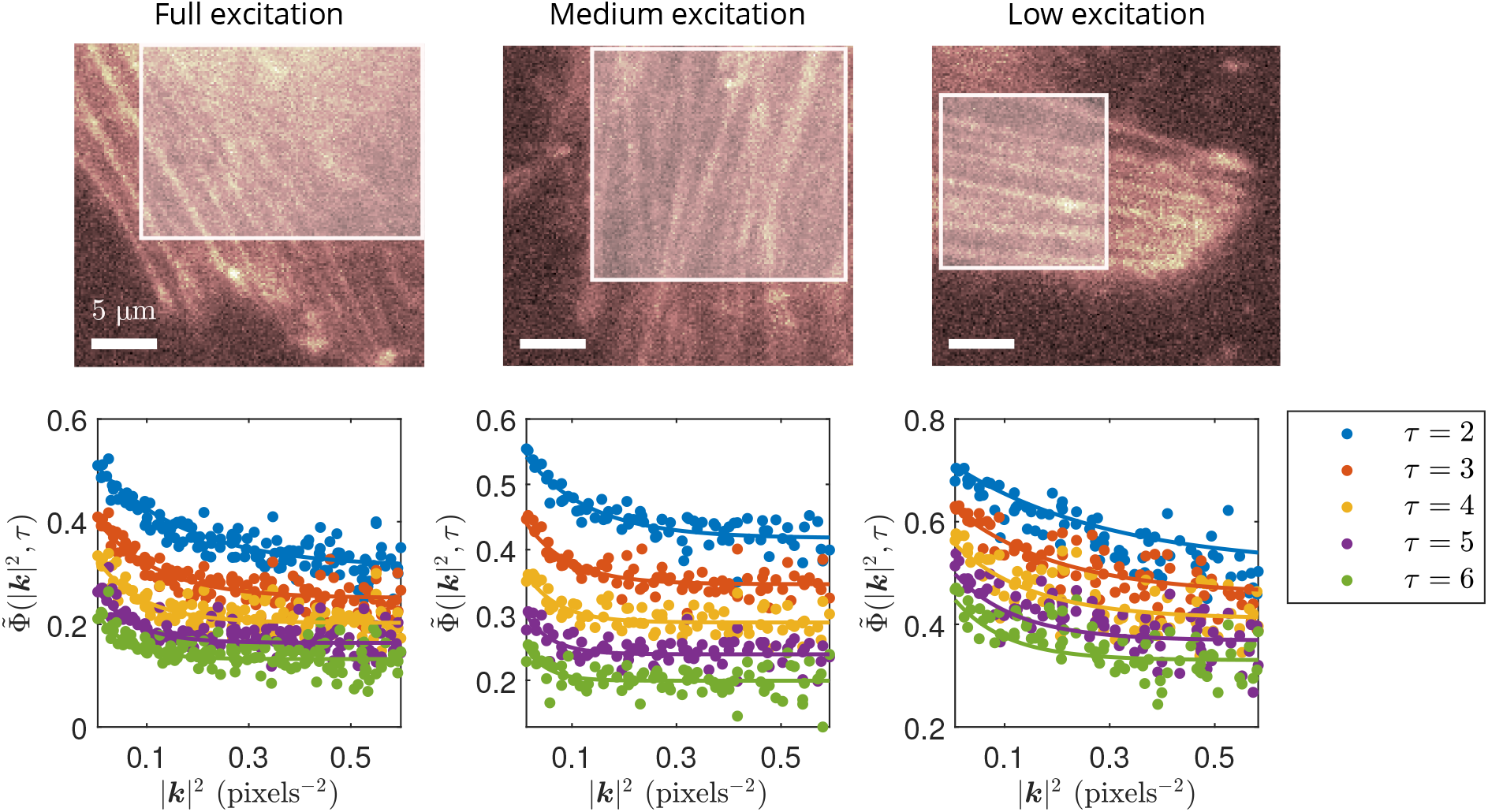
Example ACF analyses of independent HeLa cells irradiated at different excitation powers. Fluorescence images of Dronpa-C12 labeled actin in live HeLa cells, with ROI used in analyses highlighted, are shown on top part of figure. Corresponding computed ACFs from ROIs (points) and respective simultaneous fits (lines) are shown on bottom part of figure. The time lag ranges for the fits varied from *τ* = 2 to either *τ* = 10 or *τ* = 20. Time-windowing was done with *T*_w_ = 100 frames. Pixel size: 177.78 nm. Frame time: 10 ms. Image series length: 50 s.

**Table III:**
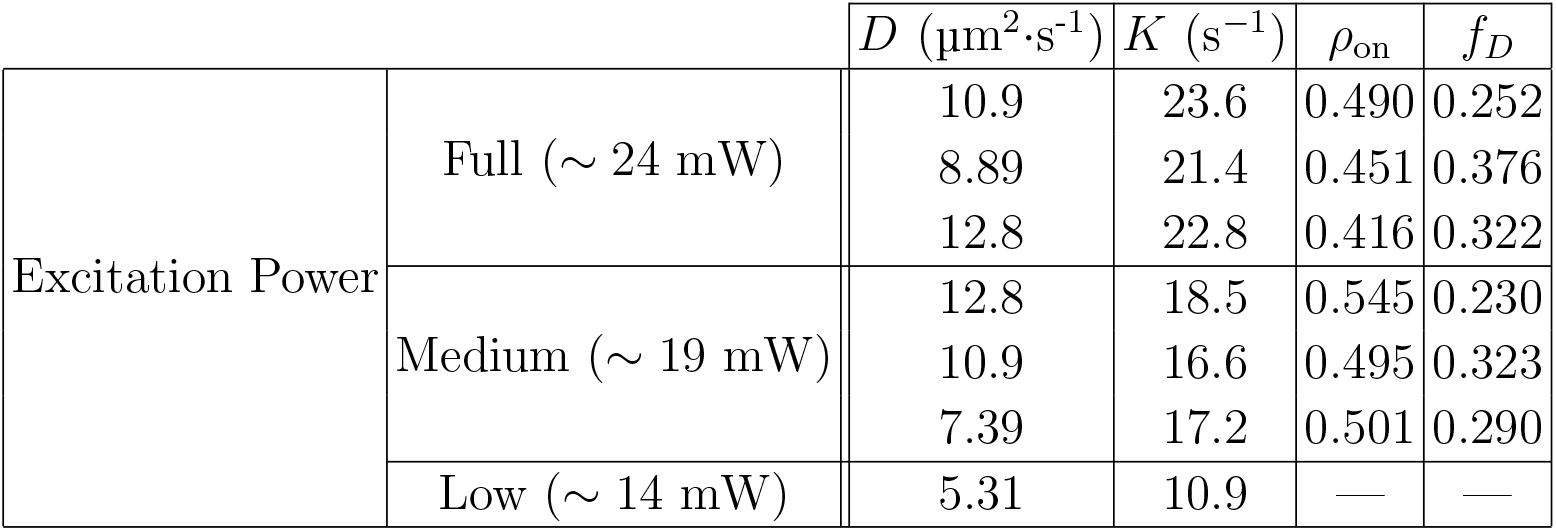
Fit parameters measured from independent HeLa cells at different excitation powers.

| | | $D$ ($\mu\text{m}^2 \cdot \text{s}^{-1}$) | $K$ ($\text{s}^{-1}$) | $\rho_{\text{on}}$ | $f_D$ |
| --- | --- | --- | --- | --- | --- |
| Excitation Power | Full ($\sim 24$ mW) | 10.9 | 23.6 | 0.490 | 0.252 |
|  |  | 8.89 | 21.4 | 0.451 | 0.376 |
|  |  | 12.8 | 22.8 | 0.416 | 0.322 |
| | Medium ($\sim 19$ mW) | 12.8 | 18.5 | 0.545 | 0.230 |
|  |  | 10.9 | 16.6 | 0.495 | 0.323 |
|  |  | 7.39 | 17.2 | 0.501 | 0.290 |
| | Low ($\sim 14$ mW) | 5.31 | 10.9 | — | — |

Measurements of the apparent diffusion coefficient from the HeLa cells are again within the range of ~ 3–30 μm^2^·s^−1^ for G-actin diffusion in cytoplasm. Furthermore, the diffusion coefficients are about the same as we measured in the 3T3 fibroblast cell. The diffusion coefficient measured from the low power dataset, however, is lower than the ones measured at higher powers. One possible explanation for this observation is the lower excitation power would lead to lower excitation probabilities, especially outside of the focal plane. As such, the signal-to-noise ratio (SNR) may not be sufficient to detect as many of the fluorescent proteins diffusing in 3D away from the focal plane, thus reducing the measured apparent diffusion coefficient (these events might be characterized by our analysis as photoblinking, for instance).

We also observed that the sum of the photoblinking rates, *K*, and on-time fraction, *ρ*_on_, consistently increased with decreasing excitation intensity. Computing the mean on-time residency from Table III we obtained *t*_on_ = 81 ± 3 ms at full excitation power and *t*_on_ = 118 ± 1 ms at medium excitation power (note, we use the convention *k*_off_ ≡ 1/*t*_on_). We also calculated the mean off-time to be *t*_off_ = 99 ± 6 ms at full excitation and *t*_off_ = 113 ± 7 ms at medium excitation. The increasing on-time with decreasing excitation intensity, as well as the roughly constant off-time, is characteristic behavior of any fluorophore because of the long-lived triplet state (or any similar dark state that depletes the ground singlet state).[52] This effect on the photoblinking rates as a function of excitation power was also observed in wild type Dronpa.[53]

We point out that Dronpa is expected to have a non-emissive state with a longer off-time,[53] while we found *t*_on_ ≃ *t*_off_. In fact, Habuchi et al. (2005)[53] found that wild type Dronpa has three distinct dark states, of which one is significantly longer than the others. The *t*_off_ values we measured can, therefore, depend on the residency times of multiple off-states. We make a simplifying argument to illustrate why our measurement may not be able to detect a much longer off-time. In our technique, we explicitly have *ρ*_on_ ≡ *k*_on_*/K*. If we now assume that *k*_on_ is a sum of two rates, say 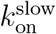 and 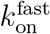, such that 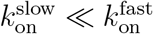, then we have 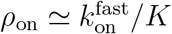. In other words, the rate corresponding to the longer time does not contribute to *ρ*_on_ and is thus not detected by our analysis.

At low power, the measured *K* is consistent with a longer on-time, assuming *t*_on_ is the only characteristic time that is a function of excitation intensity. Note we could not properly measure *ρ*_on_ and *f_D_* for the low-power data as different fitting ranges of *τ* and |***k***|^2^ would significantly affect these fitted parameters. This is, once more, likely due to the lower SNR, causing the ACF to be noisier. In general, *ρ*_on_ and *f_D_* varied most among the fitted parameters when fitting the ACF over different ranges.

We also found a mean value for the diffusing fraction of *f_D_* = 0.30 ± 0.02. Blikstad and Carlsson (1982)[54] have previously reported values of unpolymerized actin measured from HeLa cell homogenates between 35–45%.

Note the time-integrated 3D diffusion model did not fit our HeLa cell data well. This is possibly due to the non-negligible dead time (~ 1 ms) of the EMCCD camera detector relative to the shorter frame times used for imaging this data (10 ms). Instead of accounting for this effect in our model, we found that simply excluding the first time-lag from our analyses gave reasonable fits.

Different rows are ROI analyses of independent cells. Fits for *ρ*_on_ and *f_D_* at low power were omitted due to inconsistent fitted values when using different *τ* and |***k***|^2^ fitting ranges.

## CONCLUSION

We have presented an extended kICS fluorescence fluctuation rapid analysis method that simultaneously fits for the diffusion coefficient, photoblinking rates and fraction of diffusing particles from a fluorescence image series. This is done independently from any other parameters. Unlike other image correlation techniques, our current approach can be applied to regions with non-uniform fluorophore distributions, including complex cellular morphologies. This enables us to increase spatial sampling across areas of the cell, which improves the statistical precision of the ACF and extends the dynamic range for transport coefficient measurement. Furthermore, we have shown through physically realistic simulations that we can obtain accurate fit results in the presence of photobleaching, without having to consider its effects. We also demonstrated that our method can measure an apparent diffusion coefficient of Dronpa-C12 labeled actin in live NIH/3T3 and HeLa cell data that is consistent with previous literature values. We further observed that the fitted photoblinking parameters, measured from several independent HeLa cells, gave the expected trend as a function of excitation power. Lastly, our reported values for the diffusing fractions in both 3T3 and HeLa cells agree well with literature values. We anticipate that our technique will be useful in the study of dynamics in super-resolution, due to its ability to analyze more intricate systems than previous image correlation methods.

In the future, we plan to apply our method to measure biomolecular binding kinetics since photoblinking and mean-field binding/unbinding are virtually analogous processes mathematically (under certain assumptions). Another potential application could be to use the measured photoblinking rates as probes for sensing changes in a cellular environment.

## MATERIALS AND METHODS

### Live cell imaging

The NIH/3T3 fibroblast and Hela cells were transfected with plasmids containing either the Dronpa-C12 *β*-actin fusion protein or with Lipofectamine 2000 using the standard protocol. Prior to imaging, cell culture media was gently replaced into 1x PBS buffer warmed to 37°C. The cells with prominent actin stress fibers were empirically identified for imaging. The NIH/3T3 fibroblast cell data was imaged with acquisition time of 50 ms per frame and we observed slow detaching of its focal adhesion sites during the imaging time course. The same data appeared in previous work.[44] The HeLa cells were imaged at 10 ms per frame under excitation powers of ~ 14, ~ 19 and ~ 24 mW.

Imaging was performed with an inverted fluorescence microscope using widefield imaging mode (Nikon Eclipse Ti, Tokyo) equipped with an EMCCD camera (Andor iXon, model no. DU-897E-CSO-#BV) and a standard EGFP filter cube (460/60 nm band pass excitation filter, 495 nm long pass dichroic and 520/40 nm band pass emission filter). Excitation of 485/25 nm was used (cyan option, AURA light engine, ©Lumencor, Inc., Beaverton, OR, UCA). A 60x oil immersion objective (NA=1.4) was combined with an extra 1.5x magnification module integrated into the microscope body.

### Computer simulations

Simulations were created and analyzed using MATLAB R2020a on a Dell XPS 9530 (Intel(R) Core^™^ i7 @ 2.3 GHz, 16 GB RAM) running Windows 10. Simulations were also created using MATLAB R2020a on a dedicated research server (Intel(R) Core^™^ i7 @ 3.2 GHz, 64 GB RAM) running Ubuntu version 18.04.

To simulate fluorophores on filaments, we first drew angles from a normal distribution with specified mean and standard deviation. This determined the direction of each filament in the synthetic image series. Then, starting from a filament’s endpoint (randomly distributed along simulated image edges), particles were iteratively placed along the filaments with incremental distance (in pixels) drawn from a uniform distribution on (0, 1). This process was repeated until the edge of the synthetic image was reached. Each time a simulated emitter was placed, a predefined probability determined whether the position was occupied by an aggregate. Each simulated aggregate had an assigned mean number of fluorophores following a Poisson distribution, as well as a random distance from the aggregate center (mean position) following a normal distribution with a chosen standard deviation. Simulated diffusing particles were initialized randomly within the pixel grid and allowed to diffuse with periodic boundaries.

Synthetic emitters were subject to stochastic switching between on- and off-states at specified rates to simulate photoblinking. Photobleaching was assumed to be equal from the on/off-states in all simulations. Both populations of immobile and diffusing particles were simulated to have the same photoblinking and photobleaching rates.

We then convolved the simulated image series with a 2D Gaussian function (integrated over pixel dimensions) to simulate the optical PSF. To emulate the effect of the detector integration time, we split each frame into 50 “subframes”, so that a single frame was comprised of a sum of its constituent “subframes”. For more simulation details, we refer the reader to SI:Simulation details.

Synthetic pixel intensity values were assigned using the EMCCD model presented by Hirsch et al.,[55] as was previously described in Sehayek et al.[30]

The ACF was fitted to Eq. (S7) in the SI, unless otherwise stated. The fitting model assumed one diffusing and one immobile population. The global fitting of the ACF was done using the built-in Matlab object `GlobalSearch` with `fmincon` as a local solver. The fitted parameters were chosen according to the least-squares method across the specified domain of the ACF. All fitted parameters were constrained to be greater than zero, with the added conditions *ρ*_on_, *p_D_* ≤ 1. We used uniformly drawn random numbers in the interval (0, 1) as an initial guess for all fitted parameters to demonstrate the robustness of our method. In the case where the fit did not visually match the data, we repeated the fitting process until reasonable agreement was achieved. The fits always excluded the points |***k***|^2^ = 0, as they are affected by the noise in the system. The *τ* = 0 curve was also excluded from our fits, as it does not contain any useful information when using the definition in Eq. (14). Our analyses were performed on several ROIs and/or TOIs. The reported fitted parameters and their errors were then taken to be the mean and the standard error on the mean from these analyses.

### Autocorrelation computation

The autocorrelation was calculated as:

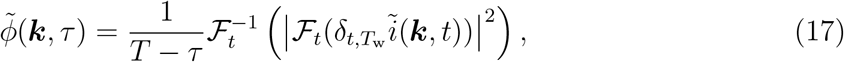

where 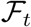 is the fast Fourier transform in time, and 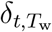 denotes the time-windowed fluctuation, as in Eq. (13); that is, at pixel (*x, y*) and frame *t*, we subtract the mean intensity of *T*_w_ subsequent frames, including frame *t* (we used Matlab `movmean` function to do this). Defining the fluctuations in this way diminishes the oscillations caused by photobleaching (see SI:Comparison with original kICS method). The choice of *T*_w_ should, ultimately, depend on the photobleaching rate. Note that the Wiener-Khinchin theorem is used in Eq. (17) to minimize autocorrelation computation time *via* Fourier (reciprocal) space calculations.

The autocorrelation in Eq. (17) was then circularly averaged. This was done by averaging all autocorrelation points with the same value of |***k***|^2^. Finally, the ACF was computed as shown in Eq. (14). To determine the “large” |***k***|^2^ offset in the denominator of the equation, a range of |***k***|^2^ values were chosen after the *τ* = 0 autocorrelation had sufficiently decayed and averaged over.

## SUPPORTING MATERIAL

- Comparison with original kICS method; autocorrelation function derivation: 2D, 3D, with time-windowed correction; simulation details; noise autocorrelation.
- kICS GitHub repository: https://github.com/ssehayek/kics-project.git
- SOFI 2.0 GitHub repository: https://github.com/xiyuyi-at-LLNL/pysofi.git

## AUTHOR CONTRIBUTIONS

- S.S. and P.W.W.: designed research, wrote the manuscript
- S.S.: developed theory for method, analyzed data, generated simulations
- X.Y. and S.W.: provided experimental data, provided SOFI analysis, contributed to method development

## ACKNOWLEDGMENTS

P.W.W. kindly acknowledges support of a Natural Science and Engineering Research Council of Canada (NSERC) Discovery Grant. S.W. was funded by the STROBE National Science Foundation Science and Technology Center, Grant No. DMR-1548924 and by the National Science Foundation, Grant No. CMI-1808766. The work from X.Y. was performed under the auspices of the U.S. Department of Energy by Lawrence Livermore National Laboratory under Contract DE-AC52-07NA27344. Release number: LLNL-JRNL-816054. Special thanks to Paul De Konick (Laval University) for providing us with an image of a branched neuron, which was used for generating simulations.

## Notes

https://github.com/ssehayek/kics-project.git

https://github.com/xiyuyi/SOFI2.0.git

### Competing Interest Statement

The authors have declared no competing interest.

### Summary of Updates

Miscellaneous edits

## References

[1] B. Huang, H. Babcock, and X. Zhuang, Breaking the Diffraction Barrier: Super-Resolution Imaging of Cells, Cell 143, 1047 (2010).

[2] E. Sezgin, Super-resolution optical microscopy for studying membrane structure and dynamics, Journal of Physics: Condensed Matter 29, 273001 (2017).

[3] M. B. Stone, S. A. Shelby, and S. L. Veatch, Super-Resolution Microscopy: Shedding Light on the Cellular Plasma Membrane, Chemical Reviews 117, 7457 (2017), pMID: 28211677, https://doi.org/10.1021/acs.chemrev.6b00716.

[4] B. O. Leung and K. C. Chou, Review of Super-Resolution Fluorescence Microscopy for Biology, Applied Spectroscopy 65, 967 (2011), pMID: 21929850, https://doi.org/10.1366/11-06398.

[5] A. Gahlmann and W. Moerner, Exploring bacterial cell biology with single-molecule tracking and super-resolution imaging, Nature Reviews Microbiology 12, 9 (2014).

[6] M. J. Rust, M. Bates, and X. Zhuang, Sub-diffraction-limit imaging by stochastic optical reconstruction microscopy (STORM), Nature methods 3, 793 (2006).

[7] E. Betzig, G. H. Patterson, R. Sougrat, O. W. Lindwasser, S. Olenych, J. S. Bonifacino, M. W. Davidson, J. Lippincott-Schwartz, and H. F. Hess, Imaging Intracellular Fluorescent Proteins at Nanometer Resolution, Science 313, 1642 (2006), https://science.sciencemag.org/content/313/5793/1642.full.pdf.

[8] T. Dertinger, R. Colyer, G. Iyer, S. Weiss, and J. Enderlein, Fast, background-free, 3D super-resolution optical fluctuation imaging (SOFI), Proceedings of the National Academy of Sciences 106, 22287 (2009), https://www.pnas.org/content/106/52/22287.full.pdf.

[9] C. Eggeling, C. Ringemann, R. Medda, G. Schwarzmann, K. Sandhoff, S. Polyakova, V. N. Belov, B. Hein, C. Von Middendorff, A. Schönle, et al., Direct observation of the nanoscale dynamics of membrane lipids in a living cell, Nature 457, 1159 (2009).

[10] A. Honigmann, V. Mueller, H. Ta, A. Schoenle, E. Sezgin, S. W. Hell, and C. Eggeling, Scanning STED-FCS reveals spatiotemporal heterogeneity of lipid interaction in the plasma membrane of living cells, Nature communications 5, 1 (2014).

[11] P. N. Hedde, R. M. Dörlich, R. Blomley, D. Gradl, E. Oppong, A. C. Cato, and G. U. Nienhaus, Stimulated emission depletion-based raster image correlation spectroscopy reveals biomolecular dynamics in live cells, Nature communications 4, 1 (2013).

[12] S. Manley, J. M. Gillette, G. H. Patterson, H. Shroff, H. F. Hess, E. Betzig, and J. Lippincott-Schwartz, High-density mapping of single-molecule trajectories with photoactivated localization microscopy, Nature methods 5, 155 (2008).

[13] F. Balzarotti, Y. Eilers, K. C. Gwosch, A. H. Gynnå, V. Westphal, F. D. Stefani, J. Elf, and S. W. Hell, Nanometer resolution imaging and tracking of fluorescent molecules with minimal photon fluxes, Science 355, 606 (2017), https://science.sciencemag.org/content/355/6325/606.full.pdf.

[14] L. Kisley, R. Brunetti, L. J. Tauzin, B. Shuang, X. Yi, A. W. Kirkeminde, D. A. Higgins, S. Weiss, and C. F. Landes, Characterization of Porous Materials by Fluorescence Correlation Spectroscopy Super-resolution Optical Fluctuation Imaging, ACS Nano 9, 9158 (2015), pMID: 26235127, https://doi.org/10.1021/acsnano.5b03430.

[15] D. Magde, E. Elson, and W. W. Webb, Thermodynamic Fluctuations in a Reacting System— Measurement by Fluorescence Correlation Spectroscopy, Phys. Rev. Lett. 29, 705 (1972).

[16] E. L. Elson and D. Magde, Fluorescence correlation spectroscopy. I. Conceptual basis and theory, Biopolymers 13, 1 (1974), https://onlinelibrary.wiley.com/doi/pdf/10.1002/bip.1974.360130102.

[17] D. Magde, E. L. Elson, and W. W. Webb, Fluorescence correlation spectroscopy. II. An experimental realization, Biopolymers 13, 29 (1974), https://onlinelibrary.wiley.com/doi/pdf/10.1002/bip.1974.360130103.

[18] J. W. Krieger, A. P. Singh, N. Bag, C. S. Garbe, T. E. Saunders, J. Langowski, and T. Wohland, Imaging fluorescence (cross-) correlation spectroscopy in live cells and organisms, Nature protocols 10, 1948 (2015).

[19] D. L. Kolin, D. Ronis, and P. W. Wiseman, k-Space Image Correlation Spectroscopy: A Method for Accurate Transport Measurements Independent of Fluorophore Photophysics, Biophysical Journal 91, 3061 (2006).

[20] D. L. Kolin and P. W. Wiseman, Advances in image correlation spectroscopy: measuring number densities, aggregation states, and dynamics of fluorescently labeled macromolecules in cells, Cell biochemistry and biophysics 49, 141 (2007).

[21] B. Hebert, S. Costantino, and P. W. Wiseman, Spatiotemporal image correlation spectroscopy (STICS) theory, verification, and application to protein velocity mapping in living CHO cells, Biophysical journal 88, 3601 (2005).

[22] M. A. Digman, C. M. Brown, P. Sengupta, P. W. Wiseman, A. R. Horwitz, and E. Gratton, Measuring Fast Dynamics in Solutions and Cells with a Laser Scanning Microscope, Biophysical Journal 89, 1317 (2005).

[23] R. Hoebe, C. Van Oven, T. W. Gadella, P. Dhonukshe, C. Van Noorden, and E. Manders, Controlled light-exposure microscopy reduces photobleaching and phototoxicity in fluorescence live-cell imaging, Nature biotechnology 25, 249 (2007).

[24] G. Donnert, C. Eggeling, and S. W. Hell, Major signal increase in fluorescence microscopy through dark-state relaxation, Nature methods 4, 81 (2007).

[25] P. P. Mondal, R. J. Gilbert, and P. T. C. So, Photobleaching reduced fluorescence correlation spectroscopy, Applied Physics Letters 97, 103704 (2010), https://doi.org/10.1063/1.3486684.

[26] L. Song, E. Hennink, I. T. Young, and H. J. Tanke, Photobleaching kinetics of fluorescein in quantitative fluorescence microscopy, Biophysical journal 68, 2588 (1995).

[27] J. Widengren and R. Rigler, Mechanisms of photobleaching investigated by fluorescence correlation spectroscopy, Bioimaging 4, 149 (1996).

[28] D. L. Kolin, S. Costantino, and P. W. Wiseman, Sampling effects, noise, and photobleaching in temporal image correlation spectroscopy, Biophysical Journal 90, 628 (2006).

[29] M. A. Digman, R. Dalal, A. F. Horwitz, and E. Gratton, Mapping the Number of Molecules and Brightness in the Laser Scanning Microscope, Biophysical Journal 94, 2320 (2008).

[30] S. Sehayek, Y. Gidi, V. Glembockyte, H. B. Brandão, P. François, G. Cosa, and P. W. Wiseman, A High-Throughput Image Correlation Method for Rapid Analysis of Fluorophore Photoblinking and Photobleaching Rates, ACS Nano 13, 11955 (2019), pMID: 31513377, https://doi.org/10.1021/acsnano.9b06033.

[31] M. Wachsmuth, W. Waldeck, and J. Langowski, Anomalous diffusion of fluorescent probes inside living cell nuclei investigated by spatially-resolved fluorescence correlation spectroscopy, Journal of Molecular Biology 298, 677 (2000).

[32] P. Sengupta, K. Garai, J. Balaji, N. Periasamy, and S. Maiti, Measuring size distribution in highly heterogeneous systems with fluorescence correlation spectroscopy, Biophysical journal 84, 1977 (2003).

[33] M. Weiss, H. Hashimoto, and T. Nilsson, Anomalous Protein Diffusion in Living Cells as Seen by Fluorescence Correlation Spectroscopy, Biophysical Journal 84, 4043 (2003).

[34] M. Weiss, M. Elsner, F. Kartberg, and T. Nilsson, Anomalous Subdiffusion Is a Measure for Cytoplasmic Crowding in Living Cells, Biophysical Journal 87, 3518 (2004).

[35] D. S. Banks and C. Fradin, Anomalous Diffusion of Proteins Due to Molecular Crowding, Biophysical Journal 89, 2960 (2005).

[36] K. Tsekouras, A. P. Siegel, R. N. Day, and S. Pressé, Inferring diffusion dynamics from fcs in heterogeneous nuclear environments, Biophysical journal 109, 7 (2015).

[37] C. M. Brown, R. B. Dalal, B. Hebert, M. A. Digman, A. R. Horwitz, and E. Gratton, Raster image correlation spectroscopy (RICS) for measuring fast protein dynamics and concentrations with a commercial laser scanning confocal microscope, Journal of Microscopy 229, 78 (2008), https://onlinelibrary.wiley.com/doi/pdf/10.1111/j.1365-2818.2007.01871.x.

[38] H. B. Brandão, H. Sangji, E. Pandžić, S. Bechstedt, G. J. Brouhard, and P. W. Wiseman, Measuring ligand–receptor binding kinetics and dynamics using k-space image correlation spectroscopy, Methods 66, 273 (2014).

[39] B. Berne and R. Pecora, Dynamic Light Scattering: With Applications to Chemistry, Biology, and Physics (Dover Publications, 2000) Chap. 5.4.

[40] W.-T. Yip, D. Hu, J. Yu, D. A. Vanden Bout, and P. F. Barbara, Classifying the Photophysical Dynamics of Single- and Multiple-Chromophoric Molecules by Single Molecule Spectroscopy, The Journal of Physical Chemistry A 102, 7564 (1998), https://doi.org/10.1021/jp981808x.

[41] E. K. L. Yeow, S. M. Melnikov, T. D. M. Bell, F. C. De Schryver, and J. Hofkens, Characterizing the Fluorescence Intermittency and Photobleaching Kinetics of Dye Molecules Immobilized on a Glass Surface, The Journal of Physical Chemistry A 110, 1726 (2006), pMID: 16451001, https://doi.org/10.1021/jp055496r.

[42] J. Widengren, U. Mets, and R. Rigler, Fluorescence correlation spectroscopy of triplet states in solution: a theoretical and experimental study, The Journal of Physical Chemistry 99, 13368 (1995), https://doi.org/10.1021/j100036a009.

[43] S. Geissbuehler, N. L. Bocchio, C. Dellagiacoma, C. Berclaz, M. Leutenegger, and T. Lasser, Mapping molecular statistics with balanced super-resolution optical fluctuation imaging (bSOFI), Optical Nanoscopy 1, 4 (2012).

[44] X. Yi, Super resolution of Optical Fluctuation Imaging 2.0 (SOFI-2.0): Towards fast super resolved imaging of live cells, Ph.D. thesis, UCLA (2017).

[45] X. Yi, S. Son, R. Ando, A. Miyawaki, and S. Weiss, Moments reconstruction and local dynamic range compression of high order superresolution optical fluctuation imaging, Biomed. Opt. Express 10, 2430 (2019).

[46] J. McGrath, Y. Tardy, C. Dewey, J. Meister, and J. Hartwig, Simultaneous Measurements of Actin Filament Turnover, Filament Fraction, and Monomer Diffusion in Endothelial Cells, Biophysical Journal 75, 2070 (1998).

[47] D. Zicha, I. M. Dobbie, M. R. Holt, J. Monypenny, D. Y. H. Soong, C. Gray, and G. A. Dunn, Rapid Actin Transport During Cell Protrusion, Science 300, 142 (2003), https://science.sciencemag.org/content/300/5616/142.full.pdf.

[48] D. McDonald, G. Carrero, C. Andrin, G. de Vries, and M. J. Hendzel, Nucleoplasmic *β*-actin exists in a dynamic equilibrium between low-mobility polymeric species and rapidly diffusing populations, Journal of Cell Biology 172, 541 (2006).

[49] T. Kiuchi, T. Nagai, K. Ohashi, and K. Mizuno, Measurements of spatiotemporal changes in G-actin concentration reveal its effect on stimulus-induced actin assembly and lamellipodium extension, Journal of Cell Biology 193, 365 (2011).

[50] I. L. Novak, B. M. Slepchenko, and A. Mogilner, Quantitative analysis of G-actin transport in motile cells, Biophysical journal 95, 1627 (2008).

[51] A. Gasilina, T. Vitali, R. Luo, X. Jian, and P. A. Randazzo, The ArfGAP ASAP1 Controls Actin Stress Fiber Organization via Its N-BAR Domain, iScience 22, 166 (2019).

[52] S. Bretschneider, C. Eggeling, and S. W. Hell, Breaking the Diffraction Barrier in Fluorescence Microscopy by Optical Shelving, Phys. Rev. Lett. 98, 218103 (2007).

[53] S. Habuchi, R. Ando, P. Dedecker, W. Verheijen, H. Mizuno, A. Miyawaki, and J. Hofkens, Reversible single-molecule photoswitching in the GFP-like fluorescent protein Dronpa, Proceedings of the National Academy of Sciences 102, 9511 (2005), https://www.pnas.org/content/102/27/9511.full.pdf.

[54] I. Blikstad and L. Carlsson, On the dynamics of the microfilament system in HeLa cells, Journal of Cell Biology 93, 122 (1982), https://rupress.org/jcb/article-pdf/93/1/122/1075730/122.pdf.

[55] M. Hirsch, R. J. Wareham, M. L. Martin-Fernandez, M. P. Hobson, and D. J. Rolfe, A Stochastic Model for Electron Multiplication Charge-Coupled Devices – From Theory to Practice, PLOS ONE 8, 1 (2013).

