## Supporting Information for "Rapid Ensemble Measurement of Protein Diffusion, Probe Blinking and Photobleaching Dynamics in the Complex Cellular Space"

Simon Sehayek

*Department of Physics, McGill University, Montreal, QC, Canada H3A 2T8*

Xiyu Yi[a]

*Department of Chemistry and Biochemistry,*
*University of California, Los Angeles 90095, USA and*
*Lawrence Livermore National Laboratory,*
*7000 East Avenue, Livermore, CA 94550, USA*

Shimon Weiss

*Department of Chemistry and Biochemistry,*
*University of California, Los Angeles 90095, USA*
*Department of Physiology, University of California, Los Angeles 90095, USA*
*California NanoSystems Institute, University of California, Los Angeles 90095, USA and*
*Department of Physics, Institute for Nanotechnology and Advanced Materials,*
*Bar-Ilan University, Ramat-Gan 52900, Israel*

Paul W. Wiseman[†]

*Department of Physics, McGill University,*
*Montreal, QC, Canada H3A 2T8 and*
*Department of Chemistry, McGill University, Montreal, QC, Canada H3A 0B8*

[a] The work from X. Yi was performed under the auspices of the U.S. Department of Energy by Lawrence Livermore National Laboratory under Contract DE-AC52-07NA27344. Release number: LLNL-JRNL-816054.

### COMPARISON WITH ORIGINAL KICS METHOD

As discussed in the main text, applying the original kICS method[1] to an image series with an immobile blinking population yields oscillations in the ACF. The original technique defined the kICS autocorrelation without the temporal fluctuations, *i.e.*,

$$\tilde{\phi}_{\text{orig}}(\boldsymbol{k},\tau) \equiv \langle \tilde{i}(\boldsymbol{k},t)\tilde{i}^*(\boldsymbol{k},t+\tau)\rangle_t. \tag{S1}$$

Using this definition, one can follow the same steps used to derive Eq. (12) in the main text to instead obtain:

$$\begin{aligned}\tilde{\phi}_{\text{orig}}(\boldsymbol{k},\tau) = q^2|\tilde{I}(\boldsymbol{k})|^2 \times \\ \Bigg\{ \underbrace{\sum_{m=n}^{N_{\text{imm}}} \langle \Theta_{m,t}\Theta_{n,t+\tau}\rangle_t + \sum_{m\neq n}^{N_{\text{imm}}} \langle \Theta_{m,t}\rangle_t \langle \Theta_{n,t+\tau}\rangle_t \exp\left(-i\boldsymbol{k}\cdot(\boldsymbol{u}_m - \boldsymbol{u}_n)\right)}_{\text{immobile}} \\ + \underbrace{\sum_{m=n}^{N_{\text{mob}}} \langle \Theta_{m,t}\Theta_{n,t+\tau}\rangle_t \left\langle \exp\left(-i\boldsymbol{k}\cdot(\boldsymbol{u}_{m,t} - \boldsymbol{u}_{n,t+\tau})\right)\right\rangle_t}_{\text{mobile}} \Bigg\} + \tilde{\phi}_\epsilon \delta_{\tau,0}.\end{aligned} \tag{S2}$$

Notice that the cross-term in this last equation (*i.e.*, the second term) is non-zero, in general. Furthermore, there is no prospect of making it zero, as was the case when introducing the time-windowed mean subtraction. Thus, the original kICS technique is affected by oscillations caused by the individual immobile particle positions. We demonstrate this effect in Fig. S1.

In part (a) of the figure below, oscillations are caused by the immobile particle positions and the presence of photobleaching. Part (b) further demonstrates that it is, in general, insufficient to define the intensity fluctuations by simply subtracting the time average, as photobleaching will still affect the ACF, in this case. Finally, part (c) shows that using an appropriate choice of time-windowed intensity fluctuations can significantly lessen the oscillatory effect.

Note we are not claiming that the extended kICS technique developed in the main text is superior to the original one. The original method allowed one to separate transport kinetics from photophysical processes in systems without an immobile blinking population

†

of fluorophores. In this work, we extended the analysis to systems with these populations and aimed to measure diffusion coefficients, as well as photophysical rates and diffusing particle fractions.

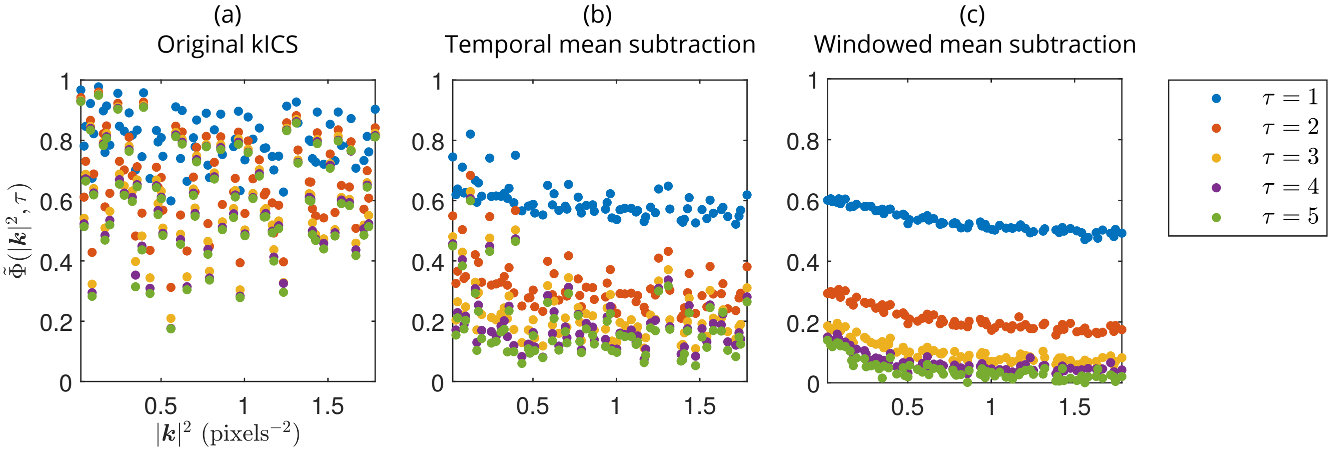


FIG. S1: Comparison of ACF (a) without temporal mean subtraction (original kICS method), (b) with temporal mean subtraction, and (c) with time-windowed mean subtraction. Time-window used in (c) is $T_{\mathrm{w}} = 200$ frames. Simulation parameters: $D = 1$ pixels$^2$·frame$^{-1}$, $K = 1$ frame$^{-1}$, $\rho_{\mathrm{on}} = 0.3$, $p_D = 0.35$ and $k_{\mathrm{p}} = 5 \times 10^{-4}$ frame$^{-1}$.

### AUTOCORRELATION FUNCTION DERIVATION

#### Diffusing and immobile populations (2D)

Here we explicitly provide the ACF for a combination of immobile and diffusing particles. We assume the fluorophores are undergoing a simple two-state, on-off photoblinking process, in the absence of photobleaching. However, we will use the expression derived here to fit for ACFs computed from systems with photobleaching. This can be a good approximation for such systems when using the time-windowed subtraction in Eq. (13) to compute the fluctuations, as discussed in the main text. In the later subsection titled "Time-windowed correction", we present a derivation that explicitly accounts for photobleaching. The fit function for the expression supplied here will also be included in the provided GitHub repository as Matlab code.

Accounting for the effect of detector time-integration in Eq. (4), the autocorrelation in

Eq. (7) is re-expressed as:

$$\tilde{\phi}(\boldsymbol{k},\tau) \equiv \begin{cases} \int_{\tau}^{\tau+1} \mathrm{d}t_2 \int_0^1 \mathrm{d}t_1 \langle \delta_t \tilde{i}(\boldsymbol{k},t_1)\delta_t \tilde{i}^*(\boldsymbol{k},t_2)\rangle_t & \tau \neq 0 \\ 2\times \int_0^1 \mathrm{d}t_2 \int_0^{t_2} \mathrm{d}t_1 \langle \delta_t \tilde{i}(\boldsymbol{k},t_1)\delta_t \tilde{i}^*(\boldsymbol{k},t_2)\rangle_t & \tau = 0 \end{cases}. \tag{S3}$$

Using the mobile component from Eq. (12), the autocorrelation for a diffusing particle is then (see Sehayek *et al.*[2] for photophysical autocorrelation details; also see Kolin *et al.*,[1] as well as Berne and Pecora[3] for the Fourier autocorrelation of diffusing particles),

$$\tilde{\phi}_{\mathrm{diff}}(Q,\tau) \equiv \begin{cases} \rho_{\mathrm{on}} e^{-Q(\tau-1)} \left( \frac{\left(1-e^{-Q}\right)^2 \rho_{\mathrm{on}}}{Q^2} + \frac{(1-\rho_{\mathrm{on}})\left(1-e^{-(Q+K)}\right)^2 e^{-K(\tau-1)}}{(Q+K)^2} \right) & \tau \neq 0 \\ 2\times \frac{1}{Q(Q+K)}\rho_{\mathrm{on}} \left( Q - \frac{Q(1-\rho_{\mathrm{on}})\left(1-e^{-(Q+K)}\right)}{Q+K} + \frac{\left(Q+e^{-Q}-1\right)K\rho_{\mathrm{on}}}{Q} - \left(1-e^{-Q}\right)\rho_{\mathrm{on}} \right) & \tau = 0 \end{cases}, \tag{S4}$$

where we define,

$$Q \equiv D|\boldsymbol{k}|^2. \tag{S5}$$

Note that in Eq. (S4), we have left out dependence on the PSF and $q$, as they are ultimately divided out by the normalization in Eq. (14).

Likewise, we obtain the autocorrelation for an immobile particle by explicitly expressing the immobile component in Eq. (12),[2]

$$\tilde{\phi}_{\mathrm{imm}}(\tau) \equiv \frac{1}{K^2}\rho_{\mathrm{on}}(1-\rho_{\mathrm{on}}) \times \begin{cases} \left(1-e^{-K}\right)^2 e^{-K(\tau-1)} & \tau \neq 0 \\ 2\times \left(e^{-K}+K-1\right) & \tau = 0 \end{cases}, \tag{S6}$$

where we again omit PSF and $q$ dependence.

It follows that the ACF, defined in Eq. (14) (including camera time-integration), for a mixture of diffusing and immobile particles is:

$$\tilde{\phi}(Q,\tau) = \frac{p_D \tilde{\phi}_{\mathrm{diff}}(Q,\tau) + (1-p_D)\tilde{\phi}_{\mathrm{imm}}(\tau)}{p_D \tilde{\phi}_{\mathrm{diff}}(Q,0) + (1-p_D)\tilde{\phi}_{\mathrm{imm}}(0)}. \tag{S7}$$

**Diffusing and immobile populations (3D)**

Here we discuss the analysis of 3D systems. We again consider the combination of immobile and diffusing populations. A full expression for the 3D ACF will be included in the provided GitHub repository as Matlab code. In the work of Kolin *et al.*,[1] it was shown that for an LSM, the 3D contribution to the kICS autocorrelation appears as a multiplying factor to its 2D counterpart. Namely, for a diffusing population, the factor is:

$$\frac{z_0^2}{4\sqrt{\pi}\sqrt{4D\tau + z_0^2}}, \tag{S8}$$

where $z_0$ is the $e^{-2}$ PSF radius in the axial direction.

Considering detector time-integration in the autocorrelation, as in Eq. (S3), the autocorrelation of a blinking, diffusing particle in 3D then has the form (in the absence of bleaching):

$$\tilde{\phi}_{\text{diff,3D}}(A, \tau) \propto \int_{\tau}^{\tau+1} \mathrm{d}t_2 \int_0^1 \mathrm{d}t_1 \frac{1}{\sqrt{4D(t_2 - t_1) + z_0^2}} e^{-A(t_2 - t_1)} \qquad (\tau \neq 0). \tag{S9}$$

This integral can be done by substituting:

$$u = \sqrt{4D(t_2 - t_1) + z_0^2}. \tag{S10}$$

Eq. (S9) is then reduced to:

$$\tilde{\phi}_{\text{diff,3D}}(B, \tau) \propto \frac{1}{2} e^{Bz_0^2} \int_{\tau}^{\tau+1} \mathrm{d}t_2 \int_{\sqrt{4D(t_2-1)+z_0^2}}^{\sqrt{4Dt_2+z_0^2}} \mathrm{d}u \, e^{-Bu^2} \qquad (\tau \neq 0), \tag{S11}$$

with

$$B \equiv A/4D. \tag{S12}$$

A similar calculation can be performed when $\tau = 0$.

For immobile populations, the 3D multiplying factor is simply $1/z_0$, as can be seen by setting $D = 0$ in Eq. (S8).

**Time-windowed correction**

Here we derive the theoretical expression for the ACF while considering the effect of the time-windowed mean subtraction. A full expression will be made available in the provided GitHub code repository. For generality, we assume the processes considered are non-stationary in time (as is the case with photobleaching, for example). Given the complexity of this expression, it is best used when the photobleaching is prominent and when Eq. (S7) cannot produce a reasonable fit to the data.

We begin by averaging the autocorrelation in Eq. (7) of the main text over all frame pairs for lag $\tau$ while using the definition of the local temporal fluctuation in Eq. (13) to obtain:

$$\tilde{\phi}(\boldsymbol{k},\tau) = \frac{1}{T-\tau}\sum_{t=0}^{T-\tau-1}\Bigg\{\tilde{g}_i(\boldsymbol{k};t,t+\tau) - \frac{1}{T_\mathrm{w}}\sum_{s=t}^{t+T_\mathrm{w}-1}\left[\tilde{g}_i(\boldsymbol{k};t,s+\tau)+\tilde{g}_i(\boldsymbol{k};s,t+\tau)-\frac{1}{T_\mathrm{w}}\sum_{s'=t}^{t+T_\mathrm{w}-1}\tilde{g}_i(\boldsymbol{k};s,s'+\tau)\right]\Bigg\}, \tag{S13}$$

where we have defined (accounting for detector time-integration in Eq. (4)):

$$\tilde{g}_i(\boldsymbol{k};u,v) \equiv \begin{cases} \int_v^{v+1}\mathrm{d}v'\int_u^{u+1}\mathrm{d}u'\,\left\langle \tilde{i}(\boldsymbol{k},u')\tilde{i}^*(\boldsymbol{k},v')\right\rangle_t & u\neq v \\ 2\times\int_u^{u+1}\mathrm{d}v'\int_u^{v'}\mathrm{d}u'\,\left\langle \tilde{i}(\boldsymbol{k},u')\tilde{i}^*(\boldsymbol{k},v')\right\rangle_t & u=v \end{cases}. \tag{S14}$$

The simplest way to carry out the sums in Eq. (S13) is to rewrite them using time-lags (see Fig. S2). We can then rewrite the first term in the square brackets of Eq. (S13) as:

$$\sum_{t=0}^{T-\tau-1}\sum_{s=t}^{t+T_\mathrm{w}-1}\tilde{g}_i(\boldsymbol{k};t,s+\tau) = \sum_{t=0}^{T-\tau-1}\sum_{\nu=0}^{T_w-1}\tilde{\phi}_i(\boldsymbol{k},\tau+\nu;t), \tag{S15}$$

where we define:

$$\tilde{\phi}_i(\boldsymbol{k},\tau;t) \equiv \tilde{g}_i(\boldsymbol{k};t,t+\tau). \tag{S16}$$

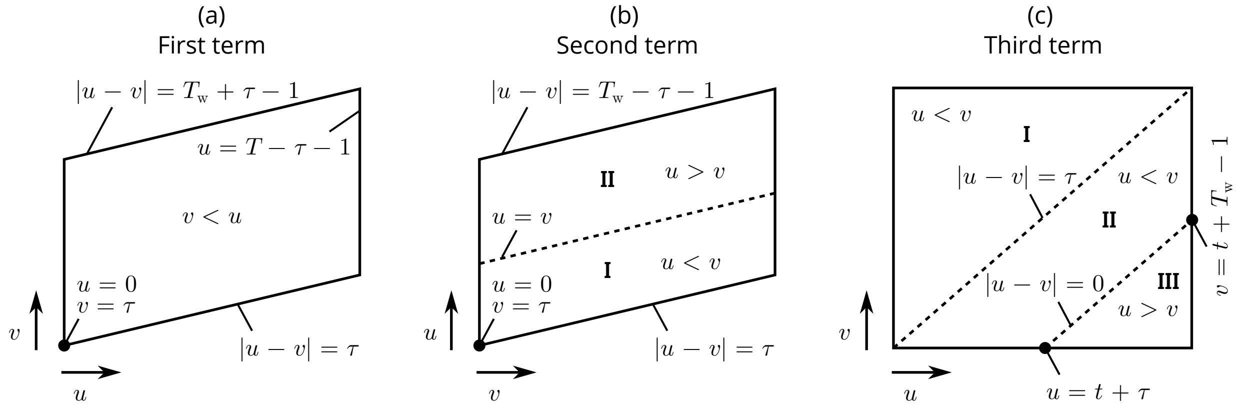


FIG. S2: Illustration of sums in square brackets of Eq. (S13). Diagonal lines within regions represent fixed time-lags, *i.e.*, constant $|u-v|$ in Eq. (S14). Sums are, therefore, simpler when carried out over and along diagonals. Time ordering of $u$ and $v$ is also shown within different subregions. Depiction of third term in (c) only shows the two innermost sums from Eq. (S13).

Using this last definition, the time-index, $t$, must follow $t \equiv \min(u, v)$, such that,

$$\tilde{g}_i(\boldsymbol{k}; u, v) \to \tilde{\phi}_i\left(\boldsymbol{k}, \tau \equiv |u - v|; t \equiv \min(u, v)\right). \tag{S17}$$

We continue to rewrite the second term (according to the subregions depicted in Fig. S2),

$$\sum_{t=0}^{T-\tau-1} \sum_{s=t}^{t+T_\mathrm{w}-1} \tilde{g}_i(\boldsymbol{k}; s, t+\tau) = \underbrace{\sum_{\nu=0}^{\tau} \sum_{t=\tau-\nu}^{T-\nu-1} \tilde{\phi}_i(\boldsymbol{k}, \nu; t)}_{\mathrm{I}} + \underbrace{\sum_{\nu=1}^{T_\mathrm{w}-\tau-1} \sum_{t=\tau}^{T-1} \tilde{\phi}_i(\boldsymbol{k}, \nu; t)}_{\mathrm{II}}. \tag{S18}$$

Finally, the third term in the square brackets of Eq. (S13) can be re-expressed as:

$$\begin{aligned} &\sum_{t=0}^{T-\tau-1} \sum_{s=t}^{t+T_\mathrm{w}-1} \sum_{s'=t}^{t+T_\mathrm{w}-1} \tilde{g}_i(\boldsymbol{k}; s, s'+\tau) = \\ &\sum_{t=0}^{T-\tau-1} \Bigg( \underbrace{\sum_{\nu=0}^{T_\mathrm{w}-1} \sum_{t'=t}^{t+T_\mathrm{w}-\nu-1} \tilde{\phi}_i(\boldsymbol{k}, \tau+\nu; t')}_{\mathrm{I}} + \underbrace{\sum_{\nu=1}^{\tau} \sum_{t'=t+\nu}^{t+T_\mathrm{w}-1} \tilde{\phi}_i(\boldsymbol{k}, \tau-\nu; t')}_{\mathrm{II}} \\ &+ \underbrace{\sum_{\nu=\tau+1}^{T_\mathrm{w}-1} \sum_{t'=t}^{t+T_\mathrm{w}-\nu-1} \tilde{\phi}_i(\boldsymbol{k}, \nu-\tau; t'+\tau)}_{\mathrm{III}} \Bigg). \end{aligned} \tag{S19}$$

Notice the number of terms in different diagonals is not constant for the third term, as was the case with the other terms.

For a mixture of immobile and diffusing populations with the same photophysical properties,[1–3]

$$\left\langle \tilde{i}(\boldsymbol{k}, u)\tilde{i}^*(\boldsymbol{k}, v)\right\rangle_t \equiv e^{-k_\mathrm{p} \max(u,v)} \rho_\mathrm{on} \left(\rho_\mathrm{on} + (1-\rho_\mathrm{on}) e^{-K|u-v|}\right) \left(N_\mathrm{imm} + N_\mathrm{diff} e^{-|\boldsymbol{k}|^2 D|u-v|}\right), \tag{S20}$$

where $k_\mathrm{p}$ is the photobleaching rate, assumed to be equal from both on- and off-states. Note the last equation assumes the cross-terms due to non-identical particles in Eq. (12) are effectively zero for reasonable choice of $T_\mathrm{w}$. We also omit the PSF and $q$ from this equation as they cancel out when using the normalization in Eq. (14).

The time-window correction to the ACF must be used when the diffusion is relatively slow, as was demonstrated in Figure 2 (C). Using the time-window correction can also allow for choosing smaller windows, which is necessary when the photobleaching is more prominent. See the main text for more details.

We compare fits with and without the time-window correction in Fig. S3 and Table S1 below. Photobleaching was not accounted for in either fit model. As we expect, the fits are more accurate when using the time-window correction. Furthermore, from this figure, one can see the effect of choosing a small time-window on the ACF at small $|\boldsymbol{k}|^2$.

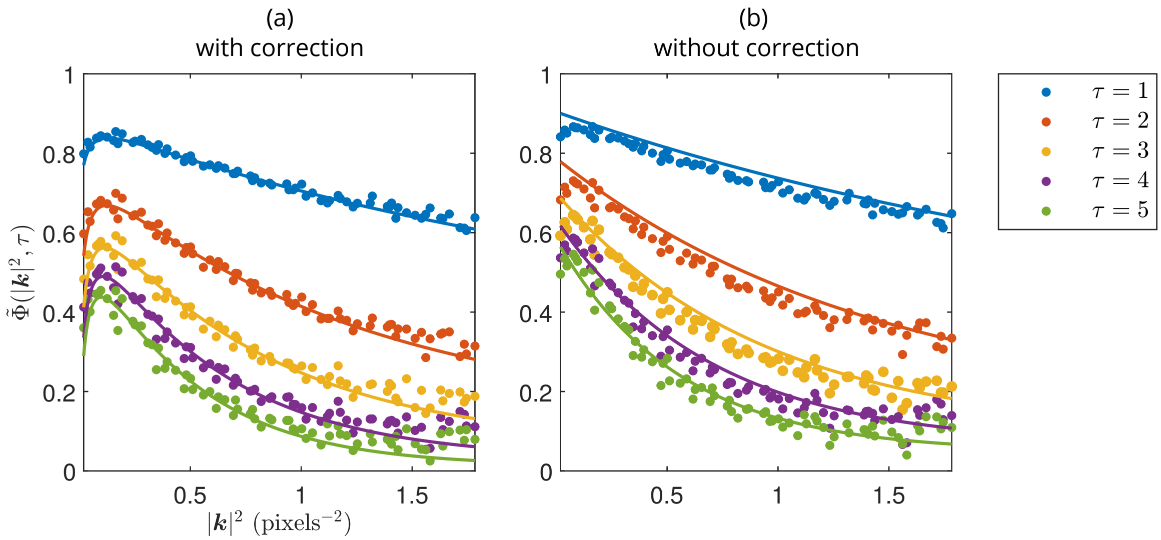


FIG. S3: Comparison of fits (a) with and (b) without time-window correction. The "bump" seen at early $|\boldsymbol{k}|^2$ is caused by the correlation of the time-window. Time windows used were (a) 50 frames and (b) 100 frames. Fit was done over first 25 time-lags, in both cases; only first 5 are shown. Photobleaching rate was set to $k_\mathrm{p} = 5 \times 10^{-4}$ frame$^{-1}$.

| | With correction | Without correction | Simulation |
|---|---|---|---|
| $D$ (pixels$^2$·frame$^{-1}$) | $0.447 \pm 0.007$ | $0.39 \pm 0.02$ | 0.5 |
| $K$ (frame$^{-1}$) | $0.60 \pm 0.02$ | $0.474 \pm 0.003$ | 0.6 |
| $\rho_\mathrm{on}$ | $0.84 \pm 0.02$ | $0.29 \pm 0.03$ | 0.833 |
| $f_D$ | $0.27 \pm 0.03$ | $0.73 \pm 0.03$ | 0.3 |

TABLE S1: Comparison of fitted and simulated parameters for fits shown in Figure S3.

Simulation was generated with $T = 2048$ frames on a $128 \times 128$ pixel grid with 4813 total particles. Fitted parameters and errors were obtained by splitting the simulation spatially into 4 equally sized and independent ROIs, and then calculating the mean and its standard error from their analyses.

### SIMULATION DETAILS

This section provides the default parameters used in our simulations (see Table S2). More details about the noise model and how we assign synthetic intensity values to the pixels in our simulations can be found in Sehayek *et al.* (2019).[2]

| Parameter description | Value |
|---|---|
| Analogue to digital conversion factor | 12 |
| Autofluorescent photon rate | 5% of mean simulated image series intensity |
| Average photon rate per molecule | $5,000$ frame$^{-1}$ |
| Clock induced charge | $5 \times 10^{-3}$ frame$^{-1}$ pixel$^{-1}$ |
| Dark noise photon rate | $8 \times 10^{-4}$ frame$^{-1}$ pixel$^{-1}$ |
| Detector quantum efficiency | 0.9 |
| EM Gain | 200 |
| Exposure time ($\tau_i$) | 0.05 s frame$^{-1}$ |
| Image dimensions | $128 \times 128$ pixels$^2$ |
| Laser $e^{-2}$ radius | $2 \times \sqrt{\text{number of pixels}}$ |
| PSF $e^{-2}$ radius | 3 pixels |
| Probability of aggregation | 0.3 |
| Mean number of monomers *per* aggregate | 2 |
| Number of filaments (where applicable) | 20 |
| Standard deviation of distance between aggregate center and monomers | 0.3 pixels |

TABLE S2: Default simulation parameters.

These parameters were used in our simulations, unless otherwise stated. Some synthetic noise parameter values are negligible, but are included for the purpose of completeness.

### NOISE AUTOCORRELATION

Here we derive an expression for the autocorrelation of the Fourier transform of the noise. Assuming $\langle \epsilon(\boldsymbol{r}, t)\rangle_t \equiv \mu_\epsilon$, it follows that:

$$\langle \tilde{\epsilon}(\boldsymbol{k}, t)\rangle_t = \int_{\mathrm{ROI}} \mathrm{d}\boldsymbol{r}\, \underbrace{\langle \epsilon(\boldsymbol{r}, t)\rangle_t}_{\mu_\epsilon} e^{-i\boldsymbol{k}\cdot\boldsymbol{r}} \overset{A_{\mathrm{ROI}}\to\infty}{\propto} \delta(\boldsymbol{k}). \tag{S21}$$

In this last equation, $A_{\mathrm{ROI}}$ denotes the area of the chosen ROI. Therefore, we have:

$$\langle \delta_t \tilde{\epsilon}(\boldsymbol{k}, t)\delta_t \tilde{\epsilon}^*(\boldsymbol{k}, t+\tau)\rangle_t = \langle \epsilon(\boldsymbol{k}, t)\tilde{\epsilon}^*(\boldsymbol{k}, t+\tau)\rangle_t, \text{ for } |\boldsymbol{k}| \neq 0. \tag{S22}$$

Furthermore, by definition of white-noise:

$$\langle \tilde{\epsilon}(\boldsymbol{r}', t)\tilde{\epsilon}^*(\boldsymbol{r}'', t+\tau)\rangle_t \equiv \sigma_\epsilon^2 \delta_{\tau,0}\delta(|\boldsymbol{r}'' - \boldsymbol{r}'|), \tag{S23}$$

where $\sigma_\epsilon^2$ is the variance of the white-noise. Using Eqs. (S22) and (S23), we obtain:

$$\begin{aligned}\langle \epsilon(\boldsymbol{k}, t)\tilde{\epsilon}^*(\boldsymbol{k}, t+\tau)\rangle_t &= \int_{\mathrm{ROI}} \mathrm{d}\boldsymbol{r}' \int_{\mathrm{ROI}} \mathrm{d}\boldsymbol{r}'' \langle \tilde{\epsilon}(\boldsymbol{r}', t)\tilde{\epsilon}^*(\boldsymbol{r}'', t+\tau)\rangle_t e^{-i\boldsymbol{k}\cdot(\boldsymbol{r}''-\boldsymbol{r}')} \\ &= \sigma^2 \delta_{\tau,0} \int_{\mathrm{ROI}} \mathrm{d}\boldsymbol{r}' \int_{\mathrm{ROI}} \mathrm{d}\boldsymbol{r}'' \delta(|\boldsymbol{r}'' - \boldsymbol{r}'|) e^{-i\boldsymbol{k}\cdot(\boldsymbol{r}''-\boldsymbol{r}')} \\ &= A_{\mathrm{ROI}} \sigma^2 \delta_{\tau,0}. \end{aligned} \tag{S24}$$


[1] D. L. Kolin, D. Ronis, and P. W. Wiseman, k-Space Image Correlation Spectroscopy: A Method for Accurate Transport Measurements Independent of Fluorophore Photophysics, Biophysical Journal **91**, 3061 (2006).

[2] S. Sehayek, Y. Gidi, V. Glembockyte, H. B. Brandão, P. François, G. Cosa, and P. W. Wiseman, A High-Throughput Image Correlation Method for Rapid Analysis of Fluorophore Photoblinking and Photobleaching Rates, ACS Nano **13**, 11955 (2019), pMID: 31513377, https://doi.org/10.1021/acsnano.9b06033.

[3] B. Berne and R. Pecora, Dynamic Light Scattering: With Applications to Chemistry, Biology, and Physics (Dover Publications, 2000) Chap. 5.4.
